# Dermal EZH2 simultaneously orchestrates dermal differentiation and epidermal proliferation during murine skin development

**DOI:** 10.1101/2020.11.01.364216

**Authors:** Venkata Thulabandu, Timothy Nehila, James W. Ferguson, Radhika P. Atit

## Abstract

Skin development and patterning is dependent on factors that regulate the stepwise differentiation of dermal fibroblasts concomitant with dermal-epidermal reciprocal signaling, two processes that are poorly understood. Here we show that dermal EZH2, the methyltransferase enzyme of the epigenetic Polycomb Repressive Complex 2 (PRC2), is a new coordinator of both these processes. Dermal EZH2 activity is present during dermal fibroblast differentiation and is required for spatially restricting Wnt/β-catenin signaling to reinforce dermal fibroblast cell fate. Later in development, dermal EZH2 regulates the differentiation to reticular dermal fibroblasts and initiation of secondary hair follicles. Embryos lacking dermal Ezh2 have elevated epidermal proliferation and differentiation that can be rescued by small molecule inhibition of retinoic acid (RA) signaling. Together, our study reveals that dermal EZH2 acts as a rheostat to control the levels of Wnt/β-catenin and RA signaling to impact fibroblast differentiation cell autonomously and epidermal keratinocyte development non-cell autonomously, respectively.

## Introduction

Differentiation of multipotent progenitors towards terminal fates is an essential component of development. Transcription factors aid in the execution of cell fate selection intrinsically by controlling the expression of fate-specific genes^1–4^. Cell-cell communication can also affect differentiation of a target cell through extrinsic juxtracrine and paracrine factors^5–7^. The presence of both intrinsic transcription factors and cell-cell communication through extrinsic factors is evident and indispensable in cell fate selection during development of many organs including skin^6–8^. However, the mechanisms that control the temporal and spatial coordination of intrinsic and extrinsic factors during cell differentiation and cell-cell communication are not clearly understood. Although *in vitro* studies have identified that epigenetic complexes may regulate coordination of cell signaling of multiple pathways^9,10^, it remains unclear as to what extent and how epigenetic complexes orchestrate coordination of cellular heterogeneity and cell-cell communication in developing tissues. Here, we use the murine skin as a model to understand the role of developmental epigenetic regulators in generating dermal fibroblast heterogeneity and dermal-epidermal communication.

The murine dermis consists of dermal fibroblasts that undergo differentiation while reciprocally signaling overlying epidermal keratinocytes to induce appendages^7,11,12^. Murine dermal fibroblasts and its interaction with the epidermal keratinocytes serve as an excellent model system to address the question of whether PRC2 can coordinate cellular differentiation both cell autonomously through intrinsic factors and non-cell autonomously through inter-cellular communication. The differentiation of dermal fibroblasts and the reciprocal signaling to the epidermal keratinocytes are integral to skin patterning, development of appendages^7,12^ and healing^13,14^. In mice, by embryonic day (E) 9.5, the dermal fibroblast (DF) precursors migrate from somites to sub-epidermal locations and differentiate into DF progenitors between E11.5 and E13.5^7,15–17^. The DF progenitors then differentiate into papillary fibroblasts progenitors (PPs), reticular fibroblasts progenitors (RPs), and hair-forming dermal papilla fibroblasts (DPs), with distinct gene expression profiles as early as E14.5^17,18^ (Fig.1A). PPs, differentiate into papillary fibroblasts, and DPs which are crucial for communicating to the epidermis and stimulating hair follicle morphogenesis and cycling^18,19^. RPs differentiate into dermal white adipose tissue (DWAT), which aids in thermal insulation^18,20^, and reticular fibroblasts, which secrete dense collagen fibers to provide elasticity to the skin^21^. It is unknown whether PRC2 controls dermal fibroblasts heterogeneity and impacts dermal signaling to the epidermis during skin development.

**Figure1:**
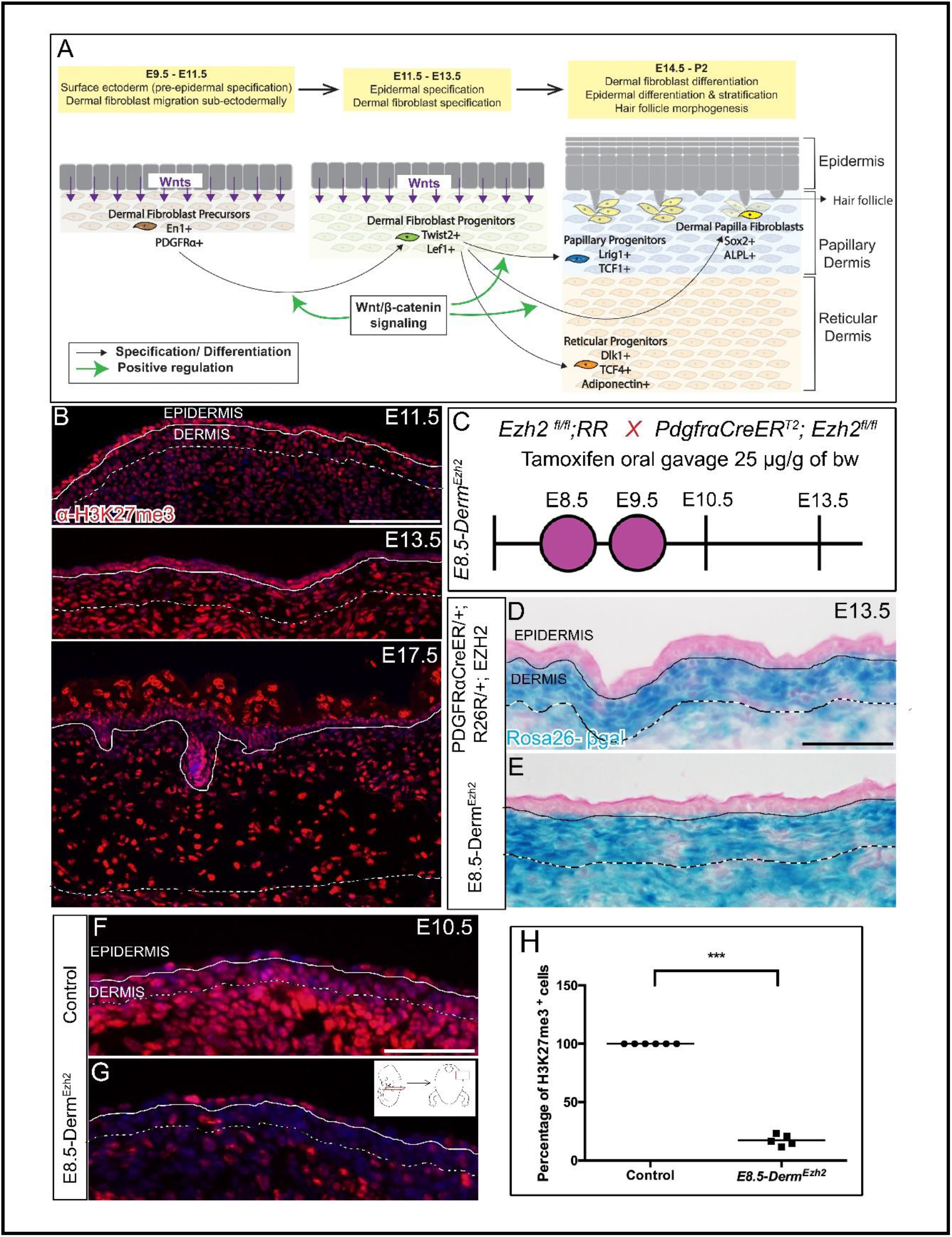
*Ezh2* is efficiently deleted in the dermal mesenchyme by PDGFRαCreER: Schematic depicting the skin development program (A). Indirect immunofluorescence of H3K27me3 with DAPI counterstain at various stages of dermal development (B). Schematic illustration of tamoxifen gavage regimen (C) utilized in the study. β-galactosidase staining (D, E) depicting the region of Cre-ER recombinase of R26R expression. Indirect immunofluorescence of H3K27me3 with DAPI counterstain at E10.5 (F, G) and its quantification (H). Solid line demarcates the epidermis from dermis and dashed line demarcates the lower limit of the dermis. Scale bar=100 microns in B and 50 microns in D-G.

Polycomb group proteins (PcG) are one of the most well-studied epigenetic complexes that regulate cell differentiation through mediation of both intrinsic and extrinsic developmental signals^22–24^. Polycomb Repressive Complex2 (PRC2) is a PcG complex consisting of three core proteins: SUZ12, EED and EZH2. EZH2 is the catalytic component that mediates the trimethylation of Lysine27 on Histone 3 (H3K27me3) resulting in gene silencing. Deleting *Ezh2* in epidermal basal keratinocytes leads to derepression of multiple late epidermal differentiation genes resulting in epidermal hyperplasia and precocious barrier acquisition^25^. Ablation of both *Ezh1* and *Ezh2* in epidermal basal keratinocytes leads to ectopic differentiation of keratinoctyes into Merkel cells mediated by an increase in *Sox2* expression^26,27^. Deleting *Ezh2* in cranial mesenchyme leads to upregulation of anti-osteogenic genes resulting in reduction of the size of skull bones^28–30^. These studies emphasize the ability of PcG to regulate differentiation cell autonomously. Besides, a previous study lends support to the non-cell autonomous regulation of PcG^31^. Ablation of *E(z),* the drosophila homologue of EZH1/2, in somatic gonad cells leads to somatic cell marker expression in drosophila germ cells in a non-cell autonomous manner^31^. These studies have demonstrated a requirement of PRC2 in regulating various intrinsic or extrinsic factors required for cell differentiation across different tissues. However, the ability of whether PRC2 coordinates various intrinsic and extrinsic factors in multipotential progenitor cells to simultaneously impact cell differentiation and cell-cell communication is not clear.

Multiple signaling pathways regulate development of DF progenitors *in vivo* during cell fate selection and differentiation. Many of the key transcription factors and signaling factors involved in DF progenitor development are regulated by PRC2 in other cell types^23,24,29^, and single gene deletions have not yielded strong dermal phenotypes. However, Wnt/β-catenin signaling regulates several transcription factors expressed between E9.5 and E11.5, such as *En1, Msx1, Msx2, Lef1, Twist1, Twist2* and is required for dermal fibroblast cell fate determination^7,15,16,32,33^.

Ablation or overexpression of *β–catenin* in DF precursors shows that canonical Wnt signaling is necessary and sufficient for specification of DF progenitors fate, respectively^7,15,16,32^. Wnt signaling expressing DF progenitors give rise to dermal condensate cells and PPs that are precursors to DP cells and are required for hair follicle initiation in embryonic mouse skin^34,35^. Wnt and BMP signaling are required to maintain DP’s ability to induce hair follicles^36^. RA signaling in DF progenitors is required to specify dermal condensate cells as shown by dermal fibroblasts restricted deletion of RA inhibitor, *Cyp26b1*^37^. Although, some of the signals governing the differentiation of DF precursors to DF progenitors have been identified, the mechanisms required to coordinate the spatio-temporal differentiation DF progenitors into PP, RP and DP lineages remain unidentified. However, whether PRC2 orchestrates these signals and factors in dermal fibroblasts is unknown.

Here, we tested the role of PRC2 in embryonic dermal development and dermal-epidermal signaling, via cell-type and temporal-specific deletion of *Ezh2* in mouse DF precursors *in vivo*. Here, we found that dermal *Ezh2* is required for differentiation of DF precursor to DF progenitors and RPs by spatially restricting Wnt/β-catenin signaling and but not for generating the other lineages. Dermal PRC2 is required for secondary hair follicle formation and dispensable for primary guard hair follicle initiation. We found that deletion of dermal *Ezh2* aberrantly enhances RA signaling and causes epidermal hyperplasia by elevating proliferation. We further demonstrate the molecular mechanism of this specific action of dermal *Ezh2.* We show that RA signaling is a key driver and transient activation of RA signaling recapitulates the phenotype and inhibition of RA signaling rescues the epidermal hyperplasia *in vivo*. Thus, dermal PRC2 directed downregulation of the RA pathway is critical for the timely activation of the epidermal proliferation and inhibition of differentiation. Altogether, these results demonstrate new and unexpected roles for dermal PRC2 to cell autonomously regulate the stepwise differentiation of dermal fibroblast cell types and non-cell autonomously regulate epidermal proliferation by maintaining critical levels of dermal Wnt/β-catenin and RA signaling.

## Results

### *Ezh2* is active throughout murine dermal development and can be efficiently ablated in dorsal dermal fibroblast precursors using PDGFRαCreER

To identify the spatio-temporal timing of PRC2 activity during dermal development, we queried PRC2-mediated H3K27me3 activity by immunostaining at various stages in mouse embryonic skin development (Fig.1A). We observed persistent expression of H3K27me3 from as early as E11.5 through E17.5 in the differentiating DFs (Fig.1B). This indicates that EZH2 is active throughout dermal development as the DF progenitors begin differentiating and signaling to the epidermis between E11.5 and E13.5.

To address the role of *Ezh2* in regulating DF differentiation, we used tamoxifen-inducible pan-mesenchymal *PDGFRαCreER*^18,38^to conditionally delete *Ezh2* and activate Rosa26 β-galactosidase (R26R) in DF precursors and broadly in the dorsal mesenchyme at E8.5 and E9.5, (E8.5-Derm^Ezh2^) (Fig.1A, 1C). As early as E10.5, we observed a 3.19-fold decrease in *Ezh2* mRNA in DF precursors in *E8.5-Derm*^*Ezh2*^ as compared to *Ezh2* heterozygous controls (Fig. S1A). The activity of *PDGFRαCreER*-mediated recombination of *R26R* is confined to the dorsal DF progenitors at E13.5 and absent in the epidermis as can be seen from β-gal staining (Fig.1D, 1E).

We visualized cells with *Ezh2* methyltransferase activity, by determining the number of H3K27me3 expressing DF precursors and progenitors. We found a 85 % decrease of H3K27me3^+^ DF precursor cells in *E8.5-Derm*^*Ezh2*^ mutants at E10.5 (Fig.1F, G, H) and 88 % decrease in DF progenitors at E13.5 (Fig.S1B, C) suggesting that *Ezh2* deletion in DF precursors is robust with our gavage regimen with *PDGFRαCreER* preceding DF differentiation.

### *Ezh2* deletion leads to upregulation of Wnt/β-catenin signaling and expansion of dermal fibroblast progenitors

Upon confirming deletion of *Ezh2* in DF precursors, we next sought to determine the effect of *Ezh2* deletion on DF precursor differentiation into DF progenitors and the signals underlying dermal fibroblast fate selection (Fig. 1A, 2A’, 2A”). The expression domain of a pan-mesenchymal marker, PDGFRα, at E13.5 was comparable between control and *E8.5-Derm*^*Ezh2*^ suggesting that cells retained their mesenchymal identity (Fig. 2B, C)^18^. We next sought to determine if DF precursor differentiation is altered in *E8.5-Derm*^*Ezh2*^ mutants. We examined the protein expression of DF progenitor markers, TWIST2 (DERMO1) and LEF1, at E12.5^7,15,16,32^. Compared to the control, the expression domain of TWIST2 and LEF1 proteins was expanded with an 84 % increase in TWIST2^+^ cells and 40 % increase in LEF1^+^ cells in the deeper layers of the *E8.5-Derm*^*Ezh2*^ dorsal mesenchyme (Fig. 2E-J). DF progenitor cell density and cell proliferation at E12.5; and dermal thickness at E17.5 (Fig. S2A-E) were not significantly different between controls and the *E8.5-Derm*^*Ezh2*^ mutants. This indicates that dermal-*Ezh2* deletion leads to an ectopic acquisition of deeper dorsal mesenchymal cells to DF progenitor identity.

**Figure2:**
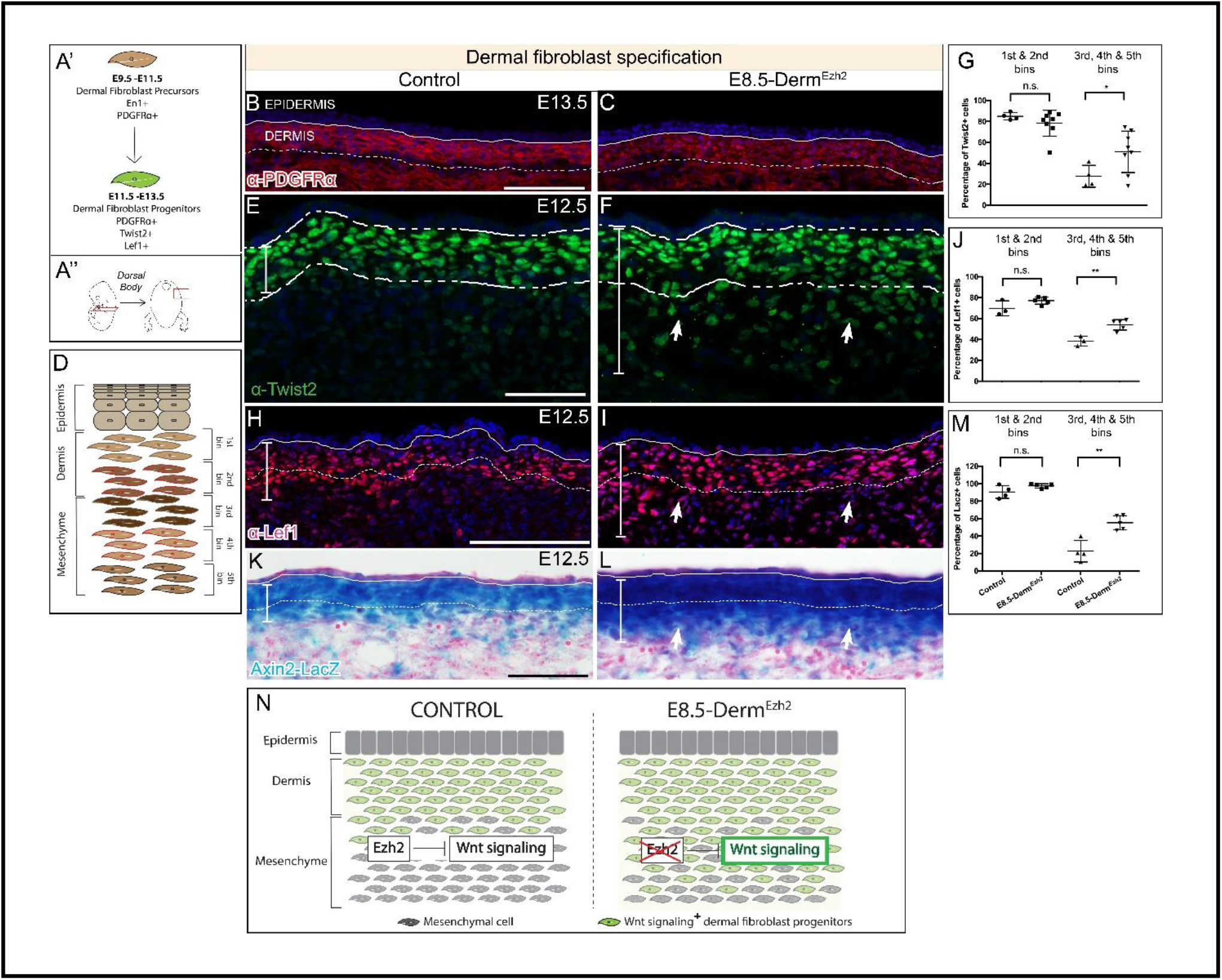
*Ezh2* restricts the specification of Wnt/ β-Catenin Signaling mediated DF progenitors to the dermis: Schematic illustration of the temporal window (A’) and region of interest (A’’). Indirect immunofluorescence with DAPI counterstain showing the expression of mesenchymal marker, PDGFRα (B, C), and dermal fibroblast progenitor markers TWIST2 (E, F), and LEF1 (H, I). *Axin2-LacZ* (β-galactosidase) staining (K, L) at E12.5. Schematic illustration of binning of dermis and dorsal mesenchyme employed to quantify cells (D). Quantification of TWIST2+ (G), LEF1+ (J), and β-galactosidase^+^ (M) cells in both controls and mutants. A summary schematic depicting the extent of Wnt/ β-Catenin signaling domain in control and expansion in theE8.5-Derm^Ezh2^ mice (N). Solid line demarcates the epidermis from dermis and dashed line demarcates the lower limit of the dermis. Scale bar=100 microns in B, C, H, I, K, L and 50 microns in E and F.

Because Wnt/β-catenin signaling is known to be involved in cell fate decisions and both TWIST2 and LEF1 are among its targets, we further investigated whether there was an upregulation of canonical Wnt/β-catenin signaling^7,16,32^. We utilized the Wnt/β-catenin signaling reporter, *Axin2-LacZ*, to look for changes in domains of Wnt signaling following loss of *Ezh2*^41,42^. β-gal staining revealed an increase in the size of the domain of *Axin2-LacZ* expression in *E8.5-Derm*^*Ezh2*^ at E12.5. We identified a 144 % increase in ectopic *Axin2-LacZ*^*+*^ cells in the deeper layers of the *E8.5-Derm*^*Ezh2*^ dorsal mesenchyme indicating an expansion of elevated canonical Wnt/β-catenin signaling (Figure 2K-M). Thus, the increase in TWIST2^+^ and LEF1^+^ DF progenitors results from an expansion of Wnt/β-catenin signaling domain mediated differentiation of dorsal mesenchymal cells underlying the dermis. These data reveal a new role for dermal PRC2 in spatially restricting Wnt signaling activity and its target genes to DF progenitors in *vivo* during dermal specification window.

### Ablation of dermal *Ezh2* promotes differentiation of dermal fibroblast progenitors towards reticular progenitors and delays secondary dermal papilla formation

As early as E14.5, dermal fibroblast progenitors differentiate into its derivates such as PPs, RPs and DPs as seen by their distinct marker expression^17,18^ (Fig.1A). Many known signaling molecules and transcription factors that regulate dermal differentiation are targets of Wnt/β-catenin signaling^18,35,51,52^. From E12.5 to E14.5, the epidermis expresses canonical Wnt ligands which are required for Wnt/β-catenin signaling in DF progenitors and subsequently the Wnt-dependent differentiation of DF progenitors into PPS and DPs^7,12^.

Wnt/β-catenin signaling is upregulated in *E8.5-Derm*^*Ezh2*^ at E12.5 and persists until E13.5 as indicated by the relative quantity of *Lef1* and *Axin2* mRNA expression in whole skin (Fig. S2F, G, TableS1). The markers of PPs, RPs and DPs are known targets of Wnt/β-catenin signaling^7,33^ (Fig.1A). So, we next examined the effect of increased Wnt/β-catenin signaling, due to *Ezh2* ablation, on the differentiation of DF Progenitors (Fig.1A). LRIG1 and TCF1 are markers of papillary progenitors, and DLK1 and TCF4 are markers of reticular progenitors^17,18^. Histomorphometric analysis revealed that protein expression domain of LRIG1 and DLK1 was comparable between E16.5 controls and *E8.5-Derm*^*Ezh2*^ in the dermis (Fig S3A-G). At E17.5, though TCF1^+^ cells are comparable between controls and *E8.5-Derm*^*Ezh2*^ mutants, we found an increase of 35 % in TCF4^+^ cells in the *E8.5-Derm*^*Ezh2*^ mutant dermis (Fig 3D, E, F). In support of this result, we observed a 11-fold increase in the relative mRNA expression of another reticular progenitor marker, *Adiponectin* (Fig.3G). However, there was no change in the relative quantities of other PP (*Lrig1, Lef1, Prdm1*) and RP markers (*Dlk1, Col4a1*) between controls and *E8.5-Derm*^*Ezh2*^ mutants in whole skin at E16.5 (Fig.3G). These embryos fail to survive birth for post-natal analysis (data not shown). Taken together, both histomorphometric and q-PCR analyses indicate that *Ezh2* promotes differentiation of DF progenitors towards reticular progenitor fate and some markers of RP lineage are targets of dermal PRC2.

**Figure3:**
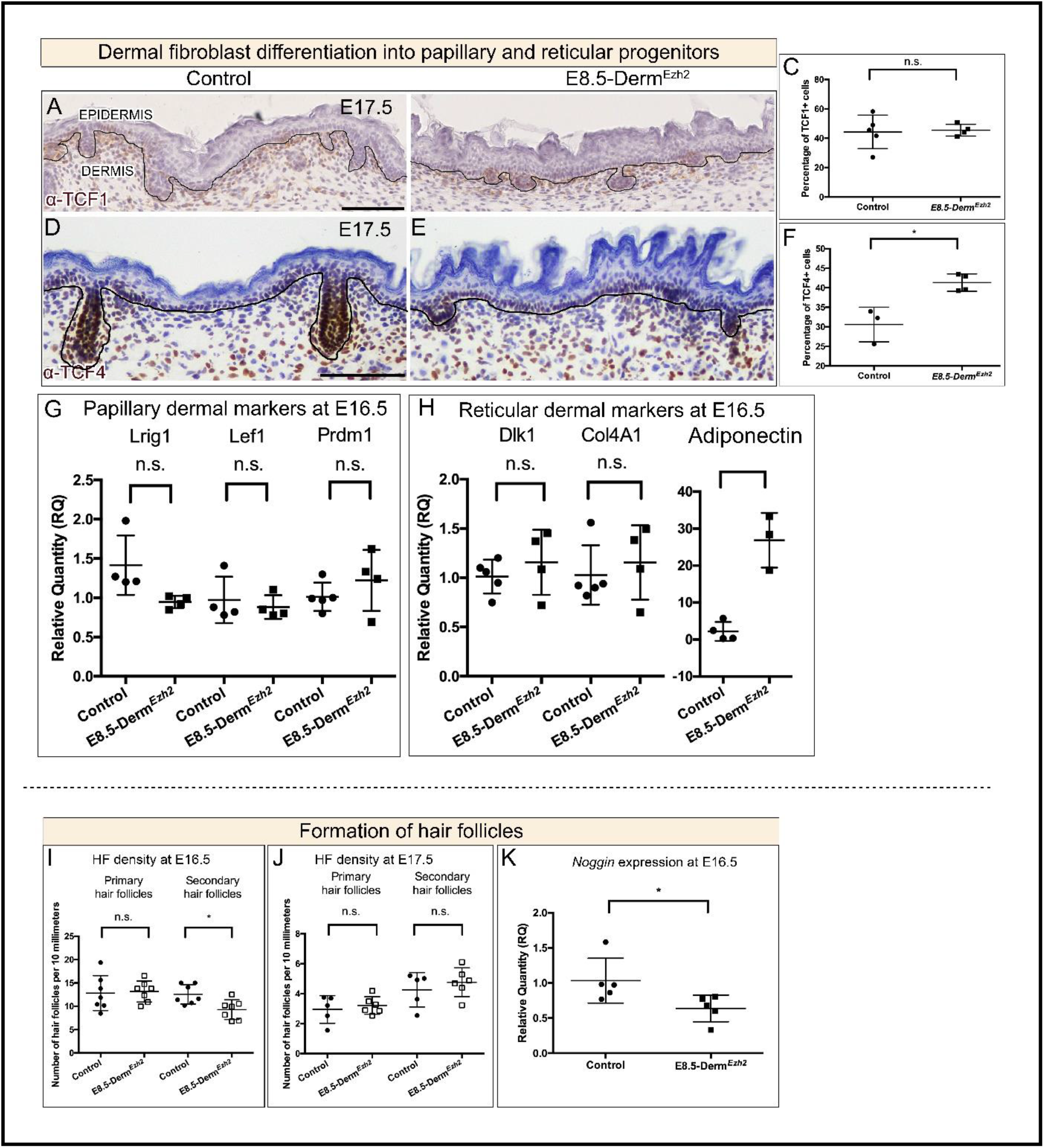
Dermal *Ezh2* ablation promotes DF progenitor differentiation towards reticular progenitors and delays secondary hair follicle initiation in *E8.5-Derm*^*Ezh2*^: Immunohistochemistry (A, B, C, D) of a PP marker, TCF1 (A, B), and an RP marker, TCF4 (D, E), with hematoxylin counterstain. Quantification of TCF1+ (C) and TCF4+ (F) cells in the dermis. The relative quantity of papillary (G) and reticular (H) progenitor markers in E16.5 whole skin between controls and *E8.5-Derm*^*Ezh2*^. Hair follicle density at E16.5 (I) and E17.5 (J). The relative quantity of secondary HF initiation gene, *Noggin* (K). Solid line demarcates the epidermis from dermis. Scale bar=100 microns in A, B, D and E.

Next, we wanted to determine the effect on differentiation of DF progenitors towards hair follicle-forming dermal papilla fibroblasts. During development, the hair follicle-forming fibroblasts make dermal condensates in the dermis which initiate hair follicles (HF) by reciprocally signaling the overlying epidermal keratinocytes to differentiate into placodal cells^5,7,12,53,54^. HF initiation occurs in three waves: primary HF at E14.5, secondary HF at E16.5 and tertiary HF at E18.5^52,55^. First, we assayed for DP differentiation by examining the expression domain of DP markers, Alkaline phosphatase^7^ and SOX2^56^ which were both comparable between controls and *E8.5-Derm*^*Ezh2*^ (Fig. S4A-D). Additionally, we surveyed the stages and number of HF in transverse sections across the entire dorsum at the forelimb level. We observed that the primary HF density was comparable between controls and *E8.5-Derm*^*Ezh2*^; however, there was a 26 % decrease in the secondary HF density at E16.5 (Fig. 3I). By E17.5, the density of primary and secondary hair follicles is comparable between controls and *E8.5-Derm*^*Ezh2*^ (Fig.3J). *Noggin* is required for secondary hair follicle initiation^57–59^. We found the relative quantities of *Noggin* mRNA was 38 % lower in *E8.5-Derm*^*Ezh2*^ (Fig.3K). These results suggest that dermal *Ezh2* ablation delays secondary hair follicle initiation possibly by impacting NOGGIN-mediated gene network without affecting differentiation of DF progenitors. Thus, dermal PRC2-EZH2 activity is dispensable for primary hair follicle initiation and is required to regulate the gene network for secondary hair follicle initiation.

### *E8.5-Derm*^*Ezh2*^ mutants exhibit epidermal hyperplasia

PRC2 activity is known to regulate various secreted factors crucial for epidermal differentiation^23,24^. We next investigated the effect of dermal *Ezh2* deletion on epidermal differentiation. We observed that *E8.5-Derm*^*Ezh2*^ mice exhibited hyperplastic epidermis throughout the dorsum skin at E16.5 (Fig4 A, B). The epidermal hyperplasia is evident as early as E13.5 (Fig. S5A-C) and besides the dorsum, the hyperplasia exists at other locations of the skin such as head skin and vibrissae skin (Fig. S5D-G). We examined the expression of KERATIN14, KERATIN10, and FILLAGRIN that are respectively expressed in basal, spinous and granular layers of the E16.5 epidermis^60,61^. We observed an expansion in the expression domain of all the three markers and it is more robust in the K10^+^ spinous layer (Fig. 4D, E, G-J). This hyperplasia is due to an increase in the number of cells: 37 % increase in basal (K14^+^) and 41 % increase in supra-basal (K10^+^ and FILAGGRIN^+^) layers (Fig.4F, K) and not due to periderm retention, which is known to delay epidermal barrier formation^62,63^. We found relative quantity of the periderm marker, Krtap13, and barrier acquisition is comparable in control and *E8.5-Derm*^*Ezh2*^ skin at E17.5 by qPCR^62^ (data not shown, Fig. S6A-D).

**Figure4:**
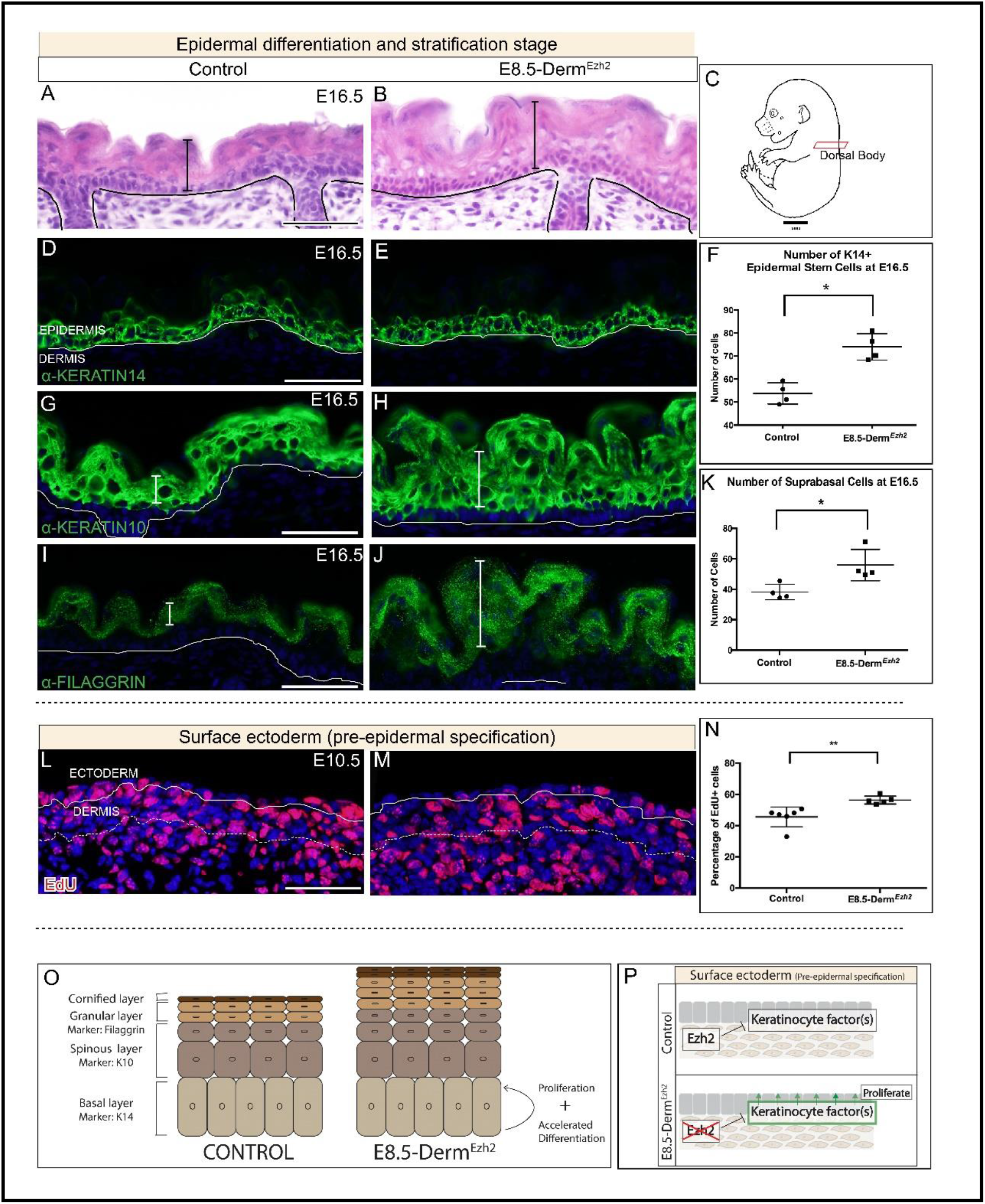
*E8.5-Derm*^*Ezh2*^ mice exhibit epidermal hyperplasia as a result of ectopic proliferation: Hematoxylin and eosin staining of dorsal body (A, B) Schematic illustration of the regions of interest (C). Indirect immunofluorescence with DAPI counterstain at E16.5 (D, E, G-H, I, J) with markers of basal keratinocytes (D, E), spinous keratinocytes (G, H) and granular keratinocytes (I, J). Percentage of basal (F) and suprabasal cells (K) in both controls and mutants. Edu staining with DAPI counterstain at E10.5 (L, M). Proliferation index of EdU^+^ basal keratinocytes in controls and mutants (N). A summary schematic depicting the effect of dermal Ezh2 deletion on the epidermis (O). Schematic with the proposed model by which dermal *Ezh2* ablation leads to epidermal hyperplasia. Solid line demarcates the epidermis from dermis and dashed line demarcates the lower limit of the dermis. Scale Bar= 50 microns.

We next sought to determine whether epidermal hyperplasia is a result of increased proliferation in *E8.5-Derm*^*Ezh2*^. Epidermal hyperplasia is first observed at E13.5, therefore we analyzed the proliferation index of the surface ectoderm at E10.5 which are the precursors of epidermal basal keratinocytes. Compared to the average of 45.53 % EdU^+^ proliferation index in controls, we found a significant increase at 56.37 % in *E8.5-Derm*^*Ezh2*^ ectoderm (Fig.4L, M, N). Lastly, we also tested whether accelerated differentiation contributes to epidermal hyperplasia. In controls, FILLAGRIN protein is expressed in the granular layer at E16.5 in the controls (Fig. 4I). However, we found FILLAGRIN protein expression was expressed by E15.5 in *E8.5-Derm*^*Ezh2*^ mice (Fig.S6E, F). We identified that hyperplasia is a stage dependent phenotype where initiation of *Ezh2* deletion at E9.5 doesn’t result in hyperplasic epidermis (data not shown). These data together suggest that dermal *Ezh2* ablation leads to elevated keratinocyte promoting factors in the skin that promote their proliferation and differentiation resulting in epidermal hyperplasia (Fig. 4O, P)

### Dermal *Ezh2* regulates cutaneous RA signaling which is sufficient to induce epidermal hyperplasia

Since dermal *Ezh2* deletion leads to epidermal hyperplasia, we next took a candidate approach to investigate if known signaling factors involved in the regulation of epidermal proliferation and differentiation are dysregulated in *E8.5-Derm*^*Ezh2*^. Administration of RA results in epidermal proliferation and subsequent epidermal thickening in human skin^64^ as seen in *E8.5-Derm*^*Ezh2*^. We queried whether RA signaling is dysregulated in *E8.5-Derm*^*Ezh2*^ skin. First, we utilized the transgenic RA signaling reporter allele, *RARE-LacZ.* β-gal staining revealed an increase in the *LacZ*^+^ cells in *E8.5-Derm*^*Ezh2*^ dermis at E10.5 (Fig.5A, B). Second, we looked at the expression of an RA target, RAR-γ, and found that there was a clear increase in RAR-γ^+^ cells in both the epidermis and dermis at E12.5 (Fig.5C, D). Both these data suggest an upregulation of RA signaling in *E8.5-Derm*^*Ezh2*^ skin.

**Figure 5:**
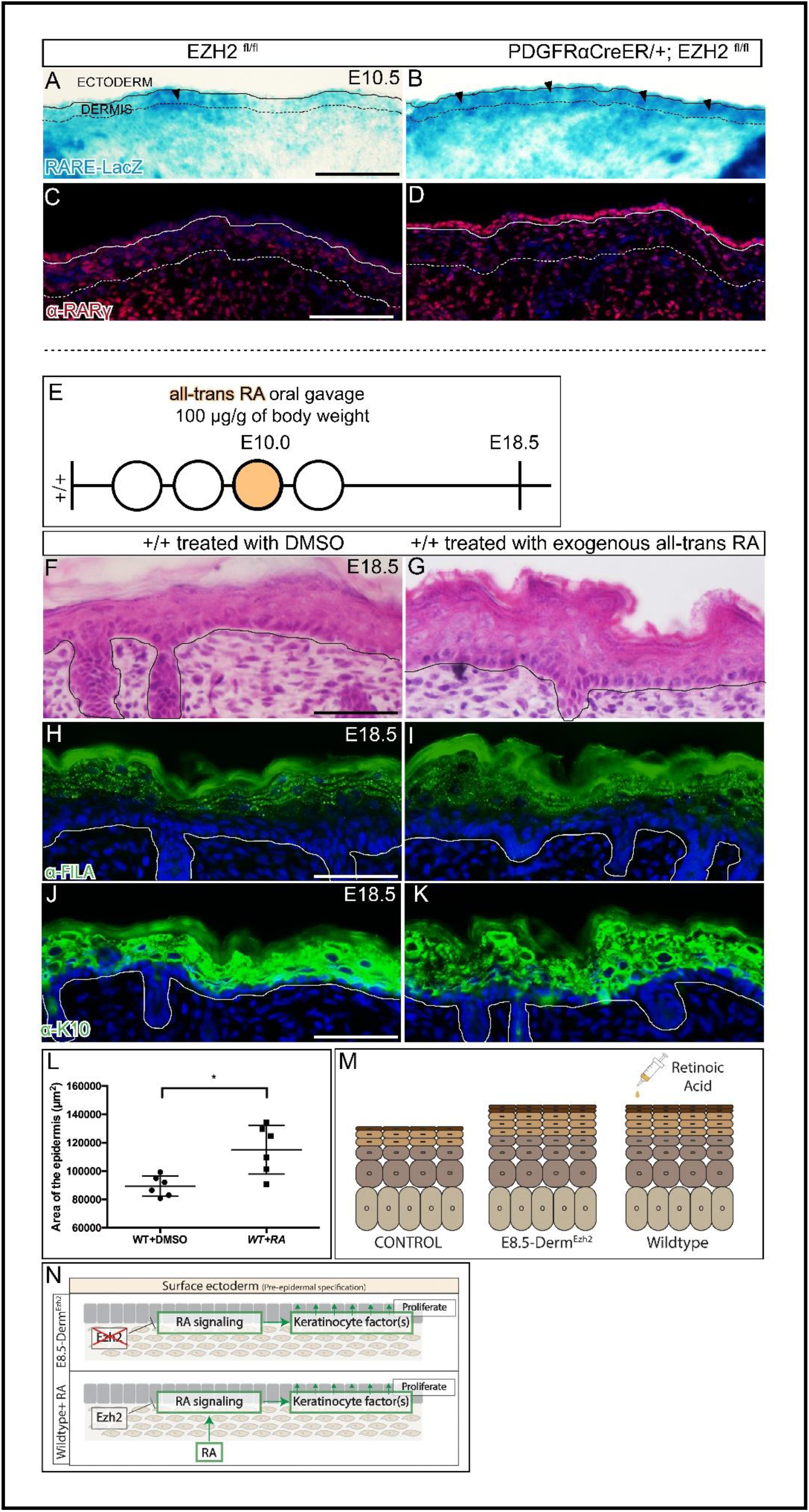
Retinoic acid is regulated by *Ezh2* and is sufficient to induce epidermal hyperplasia: β-galactosidase staining of RARE-LacZ (A, B) at E10.5. Indirect immunofluorescence with DAPI counterstain showing the expression of RA signaling target gene, RARγ (C, D) at E12.5. Schematic illustration of Retinoic Acid (E) gavage regimen. Hematoxylin and eosin staining of dorsal skin (F, G) showing the histology of the epidermis. Indirect immunofluorescence of FILAGGRIN (H, I) and KERATIN10 (J, K) with DAPI counterstain at E18.5. Comparison of epidermal area between controls and mutants in Retinoic acid treated MICE (L). A summary schematic depicting the epidermal hyperplasia phenotype (M) and the relationship between EZH2 and retinoic acid signaling (N). Solid line demarcates the epidermis from dermis and dashed line demarcates the lower limit of the dermis. Scale Bar= 50 microns.

Since retinoic acid signaling is elevated in *E8.5-Derm*^*Ezh2*^ epidermis at E10.5, we tested the functional significance of this pathway, in causing epidermal hyperplasia, using a pharmacological approach. We administered a single dose of exogenous all-trans-RA (at-RA) orally to pregnant wild-type mice carrying E10.0 embryos, the time-point at which RA signaling is dysregulated in *E8.5-Derm*^*Ezh2*^ mutants (Fig. 5E). Compared to the vehicle-treated (DMSO) controls at E18.5, we found a 28.7 % increase in epidermal area of at-RA-treated embryos (Fig. 6F, G). Similar to *E8.5-Derm*^*Ezh2*^, at-RA-treated embryos demonstrated expansion of both K10^+^ spinous and FILAGGRIN^+^ granular layers of the epidermis at E18.5 (Fig. 5H-I). These data demonstrate that a one-time administration of at-RA at E10.0 is sufficient to cause epidermal hyperplasia. In essence, dermal *Ezh2* ablation leads to upregulation of RA signaling dependent keratinocyte factors which may mediate the epidermal hyperplasia seen in *E8.5-Derm*^*Ezh2*^ mutants.

**Figure6:**
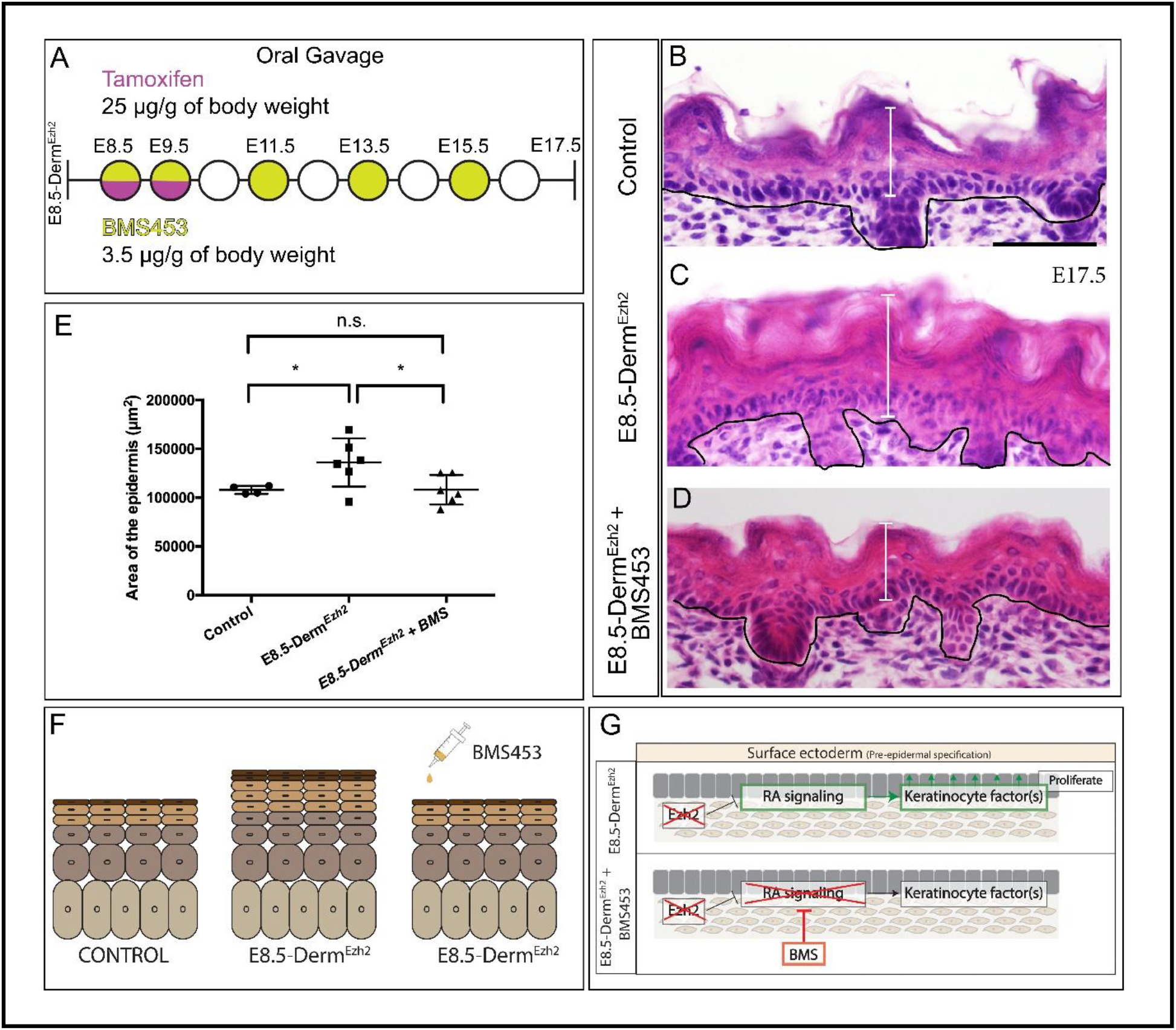
Pharmacological inhibition of Retinoic acid signaling rescues the epidermal hyperplasia in *E8.5-Derm*^*Ezh2*^: Schematic illustration BMS453 (A) gavage regimens. Hematoxylin and eosin staining of dorsal skin (B, C, D) showing the histology of the epidermis. Comparison of epidermal area between Controls, E8.5-Derm^Ezh2^ and BMS treated-E8.5-Derm^Ezh2^ mice (E). A summary schematic depicting the rescue of epidermal hyperplasia (F). The proposed model by which BMS453 is rescuing the hyperplasia in E8.5-Derm^Ezh2^ (G). Solid line demarcates the epidermal from the dermis. Scale Bar= 50 microns.

### Antagonizing RA signaling rescues epidermal hyperplasia in *E8.5-Derm*^*Ezh2*^ mice

To test that elevated RA signaling contributes to the epidermal hyperplasia observed in *E8.5-Derm*^*Ezh2*^ mice, we investigated whether antagonizing RA-signaling is sufficient to rescue epidermal hyperplasia. We orally administered BMS453, a small molecule inhibitor of RARα and RARγ^16,58^ that we validated in previous studies^29^. At 3.5μg/g body weight dosing regimen of BMS453 (Fig.6A), we rescued the epidermal area by 20.6 % as compared to the vehicle-treated (DMSO) *E8.5-Derm*^*Ezh2*^ (Fig. 6B-F). Thus, the role of dermal *Ezh2* is to ensure that RA signaling is carefully regulated to suppress keratinocyte targeted mitogenic factors and regulate normal epidermal proliferation.

In summary, our data suggest a spatio-temporal regulation of RA and Wnt/β-catenin signaling by dermal EZH2 during dermal fibroblast differentiation and epidermal-dermal signaling. We show that ablation of *Ezh2* (i) promotes Wnt/β-catenin signaling mediated differentiation of ectopic mesenchymal cells into DF progenitors (ii) promotes DF progenitor differentiation into RPs and (iii) delays secondary hair follicle formation (iv) promotes RA signaling which results in epidermal hyperplasia due to excess proliferation. Further, we rescued the epidermal hyperplasia in *Ezh2* mutants by antagonizing RA signaling with BMS453.

## Discussion

The role of PRC2 in directly regulating key developmental signaling factors and transcription factors is established in embryonic stem cells and tissue-specific stem cells^65^, but its role in coordinating spatio-temporal cell signaling events within and across tissue-specific progenitors is largely unexplored. Skin development and skin patterning with hair follicles is dependent on the coordination between the stepwise differentiation of dermal fibroblasts concomitant with dermal-epidermal reciprocal signaling. In this study, we examined the developmental role of dermal *Ezh2* during dermal fibroblast cell fate selection and reciprocal signaling with the overlying epidermis. Here we report a new action of dermal EZH2 acts as a rheostat of Wnt and RA cell signaling in the dermis. PRC2-EZH2 activity is required for spatially restricting Wnt/β-catenin signaling to reinforce DF specification and prevent lower dermal mesenchyme cells from erroneously adopting a dermal fibroblast fate. Additionally, we uncovered new roles for dermal *Ezh2* to non-cell autonomously control epidermal proliferation and differentiation by maintaining critical RA signaling levels. We found that dermal PRC2 activity is a key component of the gene regulatory network of dermal fibroblasts differentiation and dermal-epidermal signaling.

Multiple studies have demonstrated the cell autonomous regulation of PRC2-EZH2 in various cell types. In the craniofacial mesenchyme, EZH2 directly represses anti-osteogenic factors such as *Hox* genes and *Hand2* to allow for differentiation of osteoblasts into mineralized craniofacial bone^28,30^. *Ezh2* deletion in limb mesenchyme causes an increase in cell death and alteration in axis specification factors^66^. *Eed* ablation in committed cartilage progenitors, accelerates chondrocyte differentiation and premature ossification leading to a growth defect. Similarly, *Ezh2* knockout in specified calvarial bone progenitors causes premature ossification of the suture mesenchyme causing craniosynostosis of the coronal suture^29,30^. In addition, Eun et al., (2015)^31^ highlights the non-cell autonomous regulation of E(z), a homolog of EZH1/2 in drosophila, where ablation of *E(z)* in somatic gonad cells leads to somatic cell marker expression in germ cells. Although these studies demonstrated that EZH2 can exhibit cell and non-cell autonomous regulation in different contexts, they do not demonstrate whether and how EZH2 can simultaneously exhibit both during development. In this study, we found that dermal PRC2-EZH2 cell autonomously regulates differentiation of dermal fibroblast heterogeneity by inhibiting intrinsic effectors of Wnt/β-catenin signaling in the mesenchyme beneath the dermis. Simultaneously, PRC2-EZH2 non-cell autonomously regulates proliferation of epidermal basal keratinocytes by suppressing RA signaling mediated extrinsic factors.

### Dermal EZH2 represses Wnt/β-catenin signaling activation in the mesenchymal cells, underlying dermis, during cell fate selection

Wnt signaling plays a key role in mammalian embryonic dermal cell fate specification and later in differentiation of dermal condensate^7,35,52^. Ectoderm Wnts are required for canonical Wnt signaling in mouse dermal fibroblasts precursors, however it is not known how the non-cell autonomous spatial restriction of Wnt signaling is achieved^7,12^. In this study we found, deletion of dermal *Ezh2* causes expansion of Wnt signaling reporter and Wnt responsive-markers that are expressed by lineage-committed dermal fibroblast progenitor cells. However, the ectopic Wnt signaling is not sufficient to increase cell density and proliferation as seen in constitutive Wnt signaling mutants^7^. Previous studies have demonstrated that Wnt/β-catenin signaling is required for dermal fibroblast cell fate^7,15,16,32^. Wnt/β-catenin signaling at E11.5 is necessary and sufficient for *Twist2/Dermo1* and *Lef1* expression in DF progenitors, respectively^7,15,16,32^. When *Ezh2*, a component of PRC2 is ablated in pre-adipocytes *in vitro* and chondrocytes *in vivo*, Wnts and its targets are ectopically de-repressed resulting in upregulation of Wnt/β-catenin signaling^23,24^. Consistent with these studies, we show that dermal-*Ezh2* ablation leads to ectopic Wnt/β-catenin signaling at E12.5 which correlates with an expansion of TWIST2^+^ and LEF1^+^ domain of DF progenitors. Future studies will focus on rescuing the dermal *Ezh2* mutants by restoring varying levels of Wnt signaling and its interaction with RA signaling using pharmacological and genetic approaches.

Both TCF1 and TCF4 are targets of Wnt signaling. It is important to note that these targets are differentially expressed depending on the level of Wnt signaling^67^. TCF1 is a Wnt^hi^ target gene and TCF4 is a Wnt^lo^ target gene. We found that *E8.5-Derm*^*Ezh2*^ mice have more TCF4^+^ cells and comparable TCF1^+^ cells with respect to controls. These data support the inference that a higher number of dermal cells are experiencing low levels of Wnt signaling ectopically. Previous studies have shown that upregulation of Wnt signaling by stabilizing β-catenin is sufficient to significantly expand the dermis. However, the dermal thickness of dermal *Ezh2* mutants does not change, indicating that the level of upregulation of Wnt signaling seen in the dermal *Ezh2* mutants is not sufficient to induce increase in dermal proliferation and thickening. This finding follows the “dimmer switch” model proposed by Nehila et al., (2020) where knocking out epigenetic repressors such as PRC2 does not completely eliminate the target activators/ repressors, instead it serves as a rheostat to modulate expression level of activators/ repressors^68^.

### Loss of *dermal Ezh2* promotes upregulation of reticular dermal markers and delays secondary hair follicle initiation

Dermal fibroblast progenitors differentiate into PPs in the upper dermis, RPs in the lower dermis, and DC under hair follicle placodes^18,35,51^. Driskell et al., (2012) showed that dermal fibroblast progenitors start to express distinct lineage markers in the dermis as early as E14.5. The markers of both PPs, RPs, and DC lineages are regulated by Wnt signaling as shown in published RNA sequencing dataset^33,51^. It is unclear from the literature on dermal Wnt signaling knockouts and constitutive Wnt signaling mutants when and how much Wnt signaling is required for differentiation of these dermal lineages and if epigenetic regulators control levels of dermal Wnt signaling^7,12^. The level of Wnt signaling elevation seen in *Ezh2* knockout mice between E16.5 and E17.5 sufficiently induced changes in reticular dermal markers without altering papillary dermal markers. This can be explained in two possible ways: (i) that the threshold for modulating Wnt signaling is lower to induce changes in reticular dermal differentiation lineage markers than other lineages and/or (ii) Wnt signaling mediated PRC2 regulation of dermal markers at E16.5 is spatially restricted just to the reticular dermis while the papillary dermal marker regulation is independent of PRC2.

Ablation of β-catenin in DF precursors leads to a complete absence of DF progenitors and DP cells^7,12^ and stabilization of β-catenin leads to ectopic DP formation resulting in an increase in the number of hair follicles^7^. In contrast to these studies, we found a delayed initiation of secondary hair follicles in dermal *Ezh2* mutants. We show that the expression of DP markers, ALP and SOX2, are comparable between controls and *E8.5-Derm*^*Ezh2*^. So, two possible explanations of this effect are (i) that *Ezh2* negatively impacts the gene regulatory network of secondary DP initiation, but not primary DP initiation or (ii) that dermal *Ezh2* non-cell autonomously suppresses epidermal keratinocyte differentiation towards secondary follicular cell fate. These explanations are consistent with previous evidence suggesting that morphogenesis of the primary and secondary hair follicles is controlled by different mechanisms ^69–71^.

### Loss of *dermal Ezh2* induces ectopic proliferation and differentiation of epidermal keratinocytes in a non-cell autonomous manner

A unique phenotype of the dermal *Ezh2* mutant is the epidermal hyperplasia due to an increase in proliferation and accelerated differentiation of basal keratinocytes in the epidermis. Our data strongly suggest that extrinsic factors from the dermis play a causal role in epidermal hyperplasia. Addition ally, we found that exogenous at-RA exposure in WT embryos *in vivo* induced epidermal hyperplasia and RA signaling inhibitor treatment to *E8.5-Derm*^*Ezh2*^ mice rescued the hyperplasia. The association of embryonic murine epidermal hyperplasia and RA signaling was present in *Cybp26b1*^*−/−*^ mutant and dermal *Cyb26b1*^*fl/del*^, a negative regulator of RA signaling^37^.

There are several dermal signaling factors controlled by *Ezh2* during skin development that can regulate epidermal proliferation and differentiation. Among these, we propose two plausible factors that may be involved in epidermal signaling thus causing RA-mediated hyperplasia. (i) Given that epidermal metabolites are solely transferred from the dermis, elevated transient RA signaling can result from the ectopic transfer of RA metabolite from dermis to the epidermis thus leading to excessive proliferation and accelerated differentiation^72–74^. (ii) PRC2 is known to epigenetically silence loci of Wnt ligands in mesenchymal derivatives, pre-adipocytes and chondrocytes^23,24^. (iii) Wnt and RA signaling crosstalk in many contexts^75,76^. Thus, it is possible that *Ezh2* ablation derepresses Wnts in the dermal mesenchyme and promotes RA-signaling effectors that further mediate keratinocyte proliferation and accelerated differentiation^77^. Future studies will investigate the identity of *Ezh2* regulated RA-sensitive mediators and understanding the mechanism of BMS453 mediate rescue of epidermal hyperplasia.

In summary, our data supports that epigenetic regulators such as PRC2-EZH2 can simultaneously regulate intrinsic and extrinsic factors through different signaling pathways. Precisely, dermal *Ezh2* regulates intrinsic factors such as TWIST2 and LEF1 via Wnt/β-catenin signaling to control DF progenitor differentiation. Aberrant activation of Wnt/β-catenin signaling occurs in dermal fibrosis and internal organ fibrosis^78,79^. Thus, elevating PRC2 activity to repress Wnt signaling may be a new therapeutic approach to abrogate skin and organ fibrosis. Further, dermal *Ezh2* directed down regulation of RA signaling mediated extrinsic factors is critical for the timely regulation of epidermal proliferation and differentiation. This approach can be employed to treat elevated RA signaling seen in psoriatic lesions^80–82^. Our data are consistent that PRC2-EZH2 is a new “dimmer switch” or rheostat. It modulates levels or spatial restriction of Wnt/β-catenin and RA signaling *in vivo* to impact cell differentiation and rate of proliferation in cell autonomous and non-autonomous ways.

## Materials and Methods

### Key resources table

| Reagent type (species) or resource | Designation | Source or reference | Identifiers | Additional information |
| --- | --- | --- | --- | --- |
| Strain, strain background (*Mus musculus*) | Pdgfra-creER<sup>TM</sup> | JAX #018280 (Rivers et al., 2008) | *Tg(Pdgfra-cre/ERT)467Dbe* | https://www.jax.org/strain/018280 |
| Strain, strain background (*Mus musculus*) | R26R | JAX stock #003309 (Soriano, 1994) | *Gt(ROSA)26Sor<sup>tm1Sor</sup>* | https://www.jax.org/strain/003309 |
| Strain, strain background (*Mus musculus*) | Ezh2<sup>fl</sup> | JAX stock #022616 (Shen et al., 2008) | *Ezh2<sup>tm2Sho</sup>* | https://www.jax.org/strain/022616 |
| Strain, strain background (*Mus musculus*) | Conductin<sup>lacZ</sup> | JAX stock #009120 (Jho et al., 2002; Yu et al., 2005) | *Axin2<sup>tm1Wbm</sup>* | https://www.jax.org/strain/009120 |
| Strain, strain background (*Mus musculus*) | RARE-hsp68LacZ | JAX stock #008477 (Rossant et al., 1991) | *Tg(RARE-Hspa1b/lacZ)12Jrt* | https://www.jax.org/strain/008477 |
| Chemical agent | Tamoxifen | Sigma Aldrich | Cat: T5648 | 10mg/ml in corn oil |
| Chemical agent | BMS453 | Cayman chemical company | Cat: 19076 | 3.5µg/µl in DMSO |
| Chemical agent | all trans-retinoic acid | Sigma Aldrich | Cat: R2625 | 100mg/ml in DMSO |
| Antibody | rabbit anti-Twist2 | Abcam | Cat: ab66031<br>RRID:AB_2211862 | IF (1:200) |
| Antibody | goat anti-PDGFRα | R&D Systems | Cat: AF1062<br>RRID:AB_2236897 | IF (1:150) |
| Antibody | rabbit anti-H3K27me3 | Cell Signaling | Cat: 4395<br>RRID:AB_11220433 | IF (1:1000) |
| Antibody | rabbit anti-Lef1 | Cell Signaling | Cat: 2230,<br>RRID:AB_823558 | IF (1:100) |
| Antibody | goat anti-Dlk1 | R&D Systems | Cat: AF1144<br>RRID:AB_2230586 | IF (1:50) |
| Antibody | goat anti-Lrig1 | R&D Systems | Cat: AF3688<br>RRID:AB_2138836 | IF (1:100) |
| Antibody | rabbit anti-TCF4 | Cell Signaling | Cat: 2569<br>RRID:AB_2199816 | IHC (1:100) |
| Antibody | rabbit anti-TCF1 | Cell Signaling | Cat: 2203<br>RRID:AB_2199302 | IHC (1:300) |
| Antibody | rabbit anti-K14 | Covance | Cat: PRB-155P<br>RRID:AB_292096 | IF (1:2000) |
| Antibody | rabbit anti-K10 | Covance | Cat: PRB-159P<br>RRID:AB_291580 | IF (1:500) |
| Antibody | rabbit anti-Filaggrin | Covance | Cat: PRB-417P<br>RRID:AB_10064149 | IF (1:500) |
| Antibody | and rabbit anti-RARγ | Cell Signaling | Cat: D3A4<br>RRID:AB_10998934 | IF (1:1000) |
| Antibody | Goat Anti-Rabbit IgG (H+L) Antibody, Alexa Fluor 594 Conjugated | Molecular Probes | Cat: A-11012<br>RRID:AB_141359 | 1:800<br>(Secondary antibody) |
| Antibody | Donkey anti-Rabbit IgG (H+L) Highly Cross-Adsorbed Secondary Antibody, Alexa Fluor 594 | Molecular Probes | Cat: A-21207<br>RRID:AB_141637 | 1:500<br>(Secondary antibody) |
| Antibody | Donkey Anti-Rat IgG (H+L) Antibody, Alexa Fluor 488 Conjugated | Molecular Probes | Cat: A-21208<br>RRID:AB_141709 | 1:800<br>(Secondary antibody) |
| Antibody | Donkey Anti-Rat IgG (H+L) Antibody, Alexa Fluor 488 Conjugated | Molecular Probes | Cat: A-21208<br>RRID:AB_141709 | 1:500<br>(Secondary antibody) |
| Commercial assay or kit | TSA Cyanine 3 System - antibody amplification kit | Perkin Elmer | Cat: NEL704A001KT<br>RRID:AB_2572409 | Kit |
| Commercial assay or kit | MOM mouse on mouse | Vector Laboratories | Cat: BMK-2202<br>RRID:AB_2336833 | Kit |
| Commercial assay or kit | Click-iT Plus EdU kit | Thermo Fisher Scientific | Cat: C10639 | Kit |
| Sequence-based reagent | *Ezh2* | Thermo Fisher Scientific | Taqman Assay<br>Mm00468460_m1 | qPCR primers<br>Mus musculus |
| Sequence-based reagent | Cyp26b1 | Thermo Fisher Scientific | Taqman Assay Mm00558507_m1 | qPCR primers Mus musculus |
| Sequence-based reagent | Twist2 | Thermo Fisher Scientific | Taqman Assay Mm00492147_m1 | qPCR primers Mus musculus |
| Sequence-based reagent | Lef1 | Thermo Fisher Scientific | Taqman Assay Mm00550265_m1 | qPCR primers Mus musculus |
| Sequence-based reagent | Twist1 | Thermo Fisher Scientific | Taqman Assay Mm00442036_m1 | qPCR primers Mus musculus |
| Sequence-based reagent | Axin2 | Thermo Fisher Scientific | Taqman Assay Mm00443610_m1 | qPCR primers Mus musculus |
| Sequence-based reagent | Suz12 | Thermo Fisher Scientific | Taqman Assay Mm01304152_m1 | qPCR primers Mus musculus |
| Sequence-based reagent | Eed | Thermo Fisher Scientific | Taqman Assay Mm00469660_m1 | qPCR primers Mus musculus |
| Sequence-based reagent | β-actin | Invitrogen | Cat:4352663 | qPCR primers Mus musculus |
| Software, algorithm | fiji-java6-20170530 | Schindelin et al., 2012 | RRID:SCR_002285 | https://fiji.sc/ |
| Software, algorithm | CellProfiler 3.1.5 | Carpenter et al., 2006 | RRID:SCR_007358 | http://cellprofiler.org |
| Software, algorithm | Photoshop CC 19.1 | Adobe | RRID:SCR_014199 | https://www.adobe.com/products/photoshop.html |
| Software, algorithm | LAS X Core 3.5.7 | Leica Application Suite X | RRID:SCR_013673) |  |
| Software, algorithm | Graphpad Prism v8.4.2 | Graphpad | SCR_002798 |  |

### Mice

The following mice were utilized for this project: PdgfrɑCreER (JAX stock #018280)^38^, Rosa26-lacz reporter (JAX stock #003309)^39^, Ezh2 floxed (Ezh2^fl/fl^) (JAX stock #022616)^40^, *Axin2*^*LacZ*^ reporter (JAX stock #009120)^41,42^, and RARE-hsp68-LacZ reporter (JAX stock # 008477)^43^. Each of these genotypes was maintained in a mixed genetic background. The males and females were time mated and the vaginal plugs were checked daily. And a plug is assigned E0.5. Tamoxifen (Sigma Aldrich T5648) dissolved at 10mg/ml in corn oil was gavaged to mice at 25μg/g of body weight on E8.5 and E9.5. For the rescue experiments, BMS453^44–46^ (Cayman Chemical Company 19076) dissolved at 3.5μg/μl in DMSO and diluted in corn oil was gavaged to mice at 3.5μg/g of body weight on E8.5, E9.5, E11.5, E13.5, and E15.5. To the wild type mice, at-retinoic acid (Sigma Aldrich R2625) dissolved at 100mg/ml in DMSO and diluted in peanut oil was gavaged to mice at 100μg/g of body weight on E9.5 and E10.5. For each experiment a minimum of 3 controls and 3 mutants were utilized. Case Western Reserve University Institutional Animal Care and Use Committee approved all animal procedures in accordance with AVMA guidelines (Protocol 2013– 0156, Animal Welfare Assurance No. A3145-01).

### Histology and stains

Cryosectioning: (i) For cryo embedding, we followed the protocol previously published^15,29^.Once embedded, the embryos were cryo-sectioned at 10 μm and 20 μm thickness for LACZ staining in the transverse plane. (ii) For paraffin embedding, the embryos were fixed in 4 % paraformaldehyde overnight, equilibrated with 70 % ethanol overnight at 4°C, and processed for paraffin embedding. These embryos were then sectioned onto charged slides at 10 μm thickness.

Histological Stains: For β-Galactosidase detection, the cryo-sectioned slides were dried for 15mins, fixed with 4 % paraformaldehyde for 5 mins and stained with X-gal compound (Amresco 0428) dissolved at 1mg/ml for 3-6hrs at 37°C as previously described^15^. For Masson’s Trichrome staining, paraffin sectioned slides were stained with Weigert Iron Hematoxylin, Biebrich Scarlet-Acid Fuchsin Solution and Aniline Blue Solution using the standard protocol. For Alkaline phosphatase staining, cryosections were rinsed in PBS for 10 mins, fixed ice-cold Acetone on ice for 10 mins, then washed in NTMT (100mM Tris, pH9.4, 100mM NaCl, 60mM MgCl2, 0.02 % Tween-20) for 10 mins at RT. Embryos were stained in 20μl/ml NBT/BCIP (Roche 11681451001) in the dark for 10 mins at RT. Slides were washed in PBS and mounted with aqueous mounting medium.

Immunostaining: For immunostaining, the slides were dried for 15 mins, washed in PBS, blocked with buffer (PBS + 1 % BSA + 0.1 % Tween20/Triton-X-100) with 10 % goat or donkey serum for 1hr. Slides were incubated with primary antibodies overnight. Species-specific fluorescent or biotinylated secondary antibodies were incubated for 1hr to stain the slides. The primary antibodies utilized are listed in the resources table.

The EdU staining was implemented using the Click-iT Plus EdU kit (Thermo Fisher Scientific, C10639). The standard protocol suggested by the manufacturer was implemented.

Species-specific secondary antibodies utilized in this study are listed in the resources table. The TSA-Kit secondary antibody system (Perkin Elmer, NEL763E001KT) was utilized with Lef1 immunostaining.

Images were taken on Olympus BX60 microscope with the Olympus DP70 digital camera using DC controller software. Confocal images were captured using Leica TCS SP8 (Leica Biosystems) using Application Suite X software (Leica Biosystems). Images were processed in Adobe Photoshop (www.adobe.com), Fiji/ImageJ^47^, and CellProfiler™ ^48^.

Barrier Assay: The embryos at E16 and E17.5 were equilibrated with a graded series of methanol and stained in 0.1 % toluidine blue (Sigma-Aldrich T3260). The embryos were then washed in 1XPBS for 24hrs before recording the pictures.

### Tissue harvesting and RT-qPCR

*Ezh2*^*fl/fl*^ (control) and *PdgfrαCreER*; *Ezh2*^*fl/fl*^ (mutant) were used to harvest skin for gene expression analyses. For the embryos staged between E10.5 and E12.5, the dorsal halves of the bodies devoid of internal organs (carcass) were harvested. And for the embryos staged between E13.5 and E17.5, the dorsal skin was collected.

#### Fluorescence Activated Cell Sorting (FACS)

Just for flow sorting experiment, *PdgfrɑCreER*; *Rosa26-tdtomato*; *Ezh2*^*fl/+*^ (heterozygous control) and *PdgfrɑCreER*; *Rosa26-tdtomato*; *Ezh2*^*fl/fl*^ (mutant) were used in place of aforementioned genotypes. The E10.5 tissue was collected directly into 0.25 % Trypsin-EDTA (Thermo Fisher Scientific 25200056), minced, and homogenized for 4 mins at 37°C. The homogenized tissue was strained with 40μm filters (Fisher Scientific 22363547) and flow sorted into dermal (*PdgfrɑCreER; Rosa26-tdtomato* positive) and epidermal (negative) fractions using FACS-ARIA-II (P65011000099) with an 85μm nozzle. Doublets and dead cells were excluded based on forward scatter, side scatter and DAPI (0.2 mg/ml) fluorescence.

Once the select tissue/cells were collected, RNA was isolated using RNeasy MinElute cleanup kit (Qiagen 74204) with DNase treatment (Qiagen 79254), and mRNA concentration was determined using Nanodrop (ND-8000-GL). Then the mRNA was converted to cDNA using the High Capacity RNA-to-cDNA Reverse Transcription Kit (Life Technologies, 4387406).

Relative quantity of mRNA expressed was determined by using 4ng of cDNA on a StepOnePlus Real-Time PCR System (Life Technologies). Commercially available Taqman probes from Invitrogen were utilized to implement q-PCR. CT values obtained were normalized to β-actin (Invitrogen 4352663) to obtain ΔCT values. Then, the ΔCT values were normalized to the average ΔCT values of the controls to obtain ΔΔCT values and relative mRNA levels^49^.

### Histomorphometrics and automated quantification of protein expression in cell

Dermal thickness: Photos of the Masson’s Trichrome stained tissue were loaded to Fiji^47^. Using the line tool, a vertical line was drawn from the base of the epidermis (i) until the start of panniculus carnosus if present in the field (ii) until the end of the region with dense collagen staining if the panniculus carnosus was absent in the field. Lines were drawn at 5 fixed points in the field and measured for their length. The average of these 5 lengths represent the dermal thickness of the field.

Hair follicle staging: The hair follicles were staged as previously published^50^.

Epidermal area: The Hematoxylin and Eosin stained photos were loaded onto Fiji^47^. Using a freeform line tool, a box was drawn around the epidermis and the area (μm^2^) of the box was obtained.

H3K27me3 immunostaining: Images were acquired on the Olympus BX-60 with DP72 camera. The first 4 layers of the dermis was manually cropped out of the photo using the Adobe Photoshop freeform pen tool. Then the resultant photo was manually counted for percent H3K27me3^+^ cells.

TWIST2^+^ cells: Images were captured by Olympus BX-60 with DP72 camera at 20X. Using the freeform pen tool in Adobe Photoshop, three parallel lines were drawn to separate the epidermis, the first 6 layers of dermis, and the second 9 layers of dermis. Percent of cells that are TWIST2^+^ were manually counted in every region.

LEF1^+^ cells: Images were captured by Leica TCS SP8 gated STED 3X at 40X. Using the freeform pen tool in Adobe Photoshop, three parallel lines were drawn to crop the epidermis, the first 6 layers of dermis, and the underlying 9 layers of dermis into separate files. Using CellProfiler™ ^48^, we used these photos to count LEF1^+^ cells: (i) The red and blue channels were split (ii) Each channel was smoothened by Gaussian filter with automatic filter size (iii) the uneven illumination in each photo was calculated with a 60 block size and corrected (iii) The noise was reduced in blue channels (Size:7, Distance:11, Cut-off distance: 0.065) and red channels (Size:7, Distance:11, Cut-off distance: 0.09) using ‘ReduceNoise’ function (iv) The number of red and blue objects were identified with adaptive Otsu three-class thresholding with an object diameter between 5 and 40.

#### *Axin-LacZ* and *Rare-LacZ* staining

The DAPI fluorescence and LacZ staining in transmitted light were captured by Leica TCS SP8 gated STED 3X at 40X. The LacZ (transmitted light) channel was inverted and converted (pseudo-colored) to Red by default CellProfiler™ settings. The pseudo colored photo was then cropped using the freeform pen tool in Adobe Photoshop to separate the epidermis, the first 6 layers of dermis and the underlying 9 layers of dermis into separate files by drawing 3 parallel lines. Then these photos were used to count the DAPI and LacZ+ cells present in them. In order to do that, (i) the cropped photos were split into separate channels (ii) the DAPI and LacZ channels were both smoothened with a Gaussian diameter of 3 and rescaled the intensity to full (iii) Then DAPI objects were identified by adaptive Otsu three-class thresholding with an object diameter between 4 and 40 (iv) The LacZ objects were identified by manual thresholding of 0.35 and segmented by watershed method with default settings. (v) The DAPI objects that did not have 35 % of their area overlapped with LacZ objects were masked (vi) The number of unmasked DAPI objects and the total DAPI objects were noted in a file.

##### DLK^+^ and LRIG1^+^ cells

The merged photos of DAPI and red channel were loaded to Fiji. The approximate size of a box that can enclose four layers of the dermis was determined by studying multiple photos. For each photo, 5 ‘region of interest’ files were created: 1 just with the epidermis and 4 non-overlapping boxes splitting the first 16 layers of the dermis. After creating these files, the fluorescence intensity in each region of interest was measured against the background. Then to count the number of cells in each region, the merged photo was manually loaded on to Fiji and (i) Splitted into RGB channels and the blue channel is selected (ii) Gaussian smoothed with a radius of 1 (iii) Thresholding of ‘Dark background’ is implemented with automatic settings (iv) segmented with watershed (v) The number of DAPI cells were identified by analyzing the particles with a size between 0.1 and infinity, and circularity between 0 and 1. Using the fluorescence intensity and number of DAPI cells in each region, the corrected fluorescence intensity was determined.

##### TCF1^+^ and TCF4^+^ cells

The brightfield photos were manually cropped to consist of just the dermis (devoid of hair follicles and other tissues). Color deconvolution plugin in Fiji was used to separate hematoxylin and DAB staining and each of the channels were saved in separate files. Then CellProfiler™ was used to count the number of hematoxylin and DAB cells by doing the following on both the files (i) The image was inverted to brighten the dark spots and vice versa (ii) The image was transformed to grayscale (iii) the noise was reduced in both DAB (Size: 7, Distance: 11, Cut-off distance: 0.075) and hematoxylin (Size:7, Distance:11, Cut-off distance: 0.05) channels using ‘ReduceNoise’ function (iv) The number of DAB+ and Hematoxylin+ objects were identified with adaptive Otsu three class thresholding with an object diameter for DAB between 5 and 30 and for hematoxylin between 5 and 40.

##### Basal and suprabasal cell counts

The merged photos of DAPI and K14 staining were loaded to Adobe Photoshop. The cells that expressed K14 (basal cells) and the cells on top of K14+ cells (suprabasal cells) were manually counted. EdU staining: At E10.5, images were taken on Olympus BX60 microscope with the Olympus DP70 digital camera. Using the freeform pen tool in Adobe Photoshop, 2 parallel lines were drawn to divide the epidermis and from the first 4 layers of dermis into separate files. And then the percent of Edu^+^ cells were manually counted.

### Statistics

All graphs were generated using Prism 6 (GraphPad Software). All the statistical analyses were carried out in Microsoft excel using the Student’s *t*-test function using an unpaired, two-tail, with unequal variance. The p-values for statistical tests in all figures are represented as: * = P < 0.05, **= P < 0.01 and ***= P < 0.001.

## Acknowledgements

Thanks to previous and current members of the Atit Lab for their input and feedback. We thank Gregg DiNuoscio and Vidhi Mendpara for genotyping animals. We thank Dr. Elena Ezhkhova, Dr. Peggy Myung, Dr. Martin Basch, and Dr. Claudia Mizutani for critical feedback. We thank Brian Zhang, Celine Opdycke and Charlotte Lo for help with manual and automated counting. We thank the SOM Light Microscopy Core Facility and the CWRU bio[box] shared instrumentation facility.

## Author Contributions

V.T. and R.P.A. conceived experiment, analyzed and interpreted data. V.T., T.N., J.F. carried out the experiments, and V.T. generated the figures.

## Supplementary Figures

**FigureS1:**
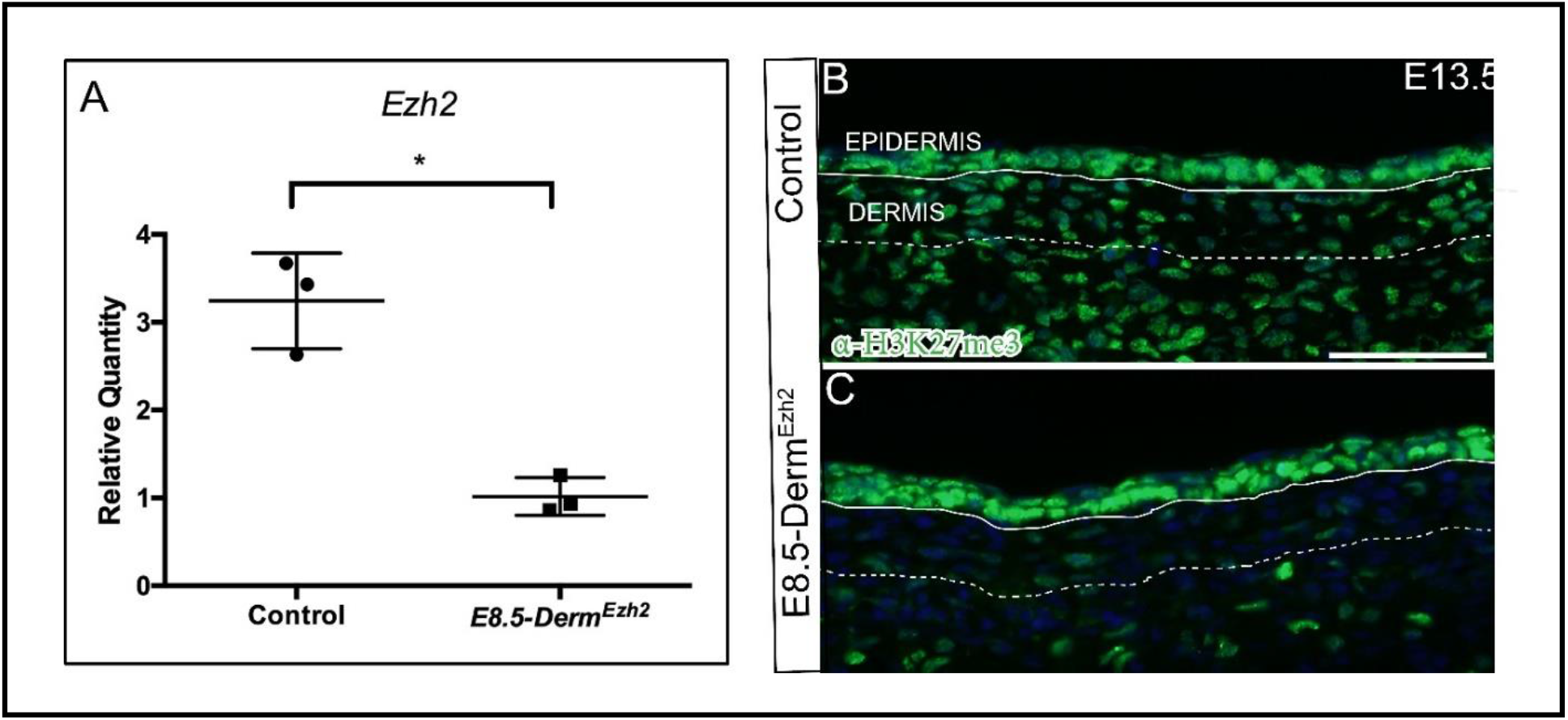
*PDGFRαCreER* efficiently deletes mesenchymal *Ezh2:* The relative quantity of *Ezh2* (A), in E10.5 flow sorted dorsal mesenchyme isolated from conditional heterozygous *E8.5-Derm*^*Ezh2 flox/+*^ controls and *E8.5-Derm*^*Ezh2*^. Indirect immunofluorescence of H3K27me3 with DAPI counterstain at E13.5 (B, C). Solid line demarcates the epidermis from dermis and dashed line demarcates the lower limit of the dermis. Scale Bar= 50 microns in B and C.

**FigureS2:**
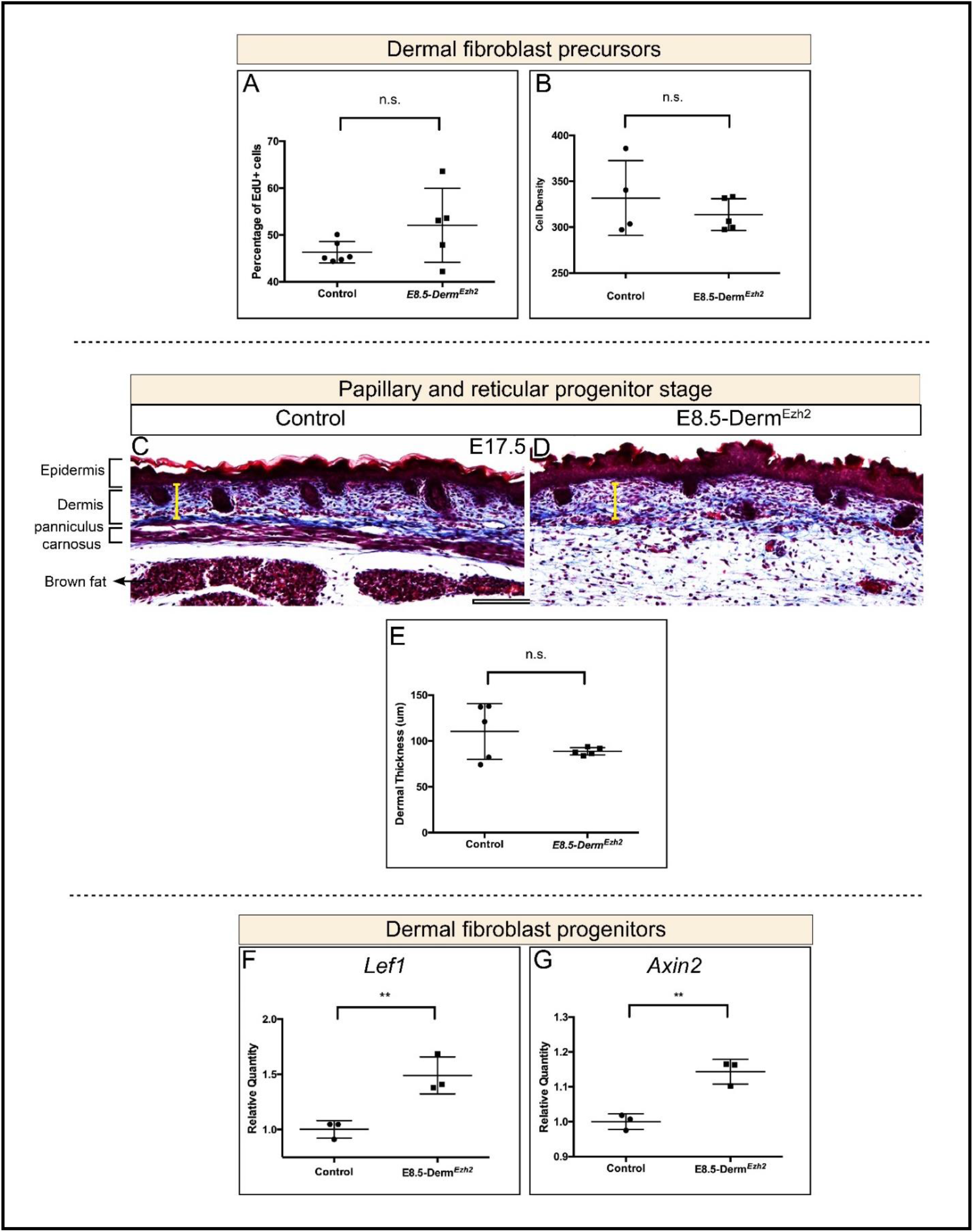
Dermal density and thickness are comparable between controls and *E8.5-Derm*^*Ezh2*^ despite the persistence of Wnt signaling upregulation till E13.5. Dermal proliferation index at E10.5 (A) and cell density (B) at E12.5. Masson’s Trichrome staining (C, D) at E17.5 showing the thickness of the dermis and its quantification (E). The relative quantity of Wnt signaling target genes, *Lef1* (F) and *Axin2* (G) in E13.5 whole skin between controls and *E8.5-Derm*^*Ezh2*^. Scale Bar= 200 microns in C and D.

**Figure S3:**
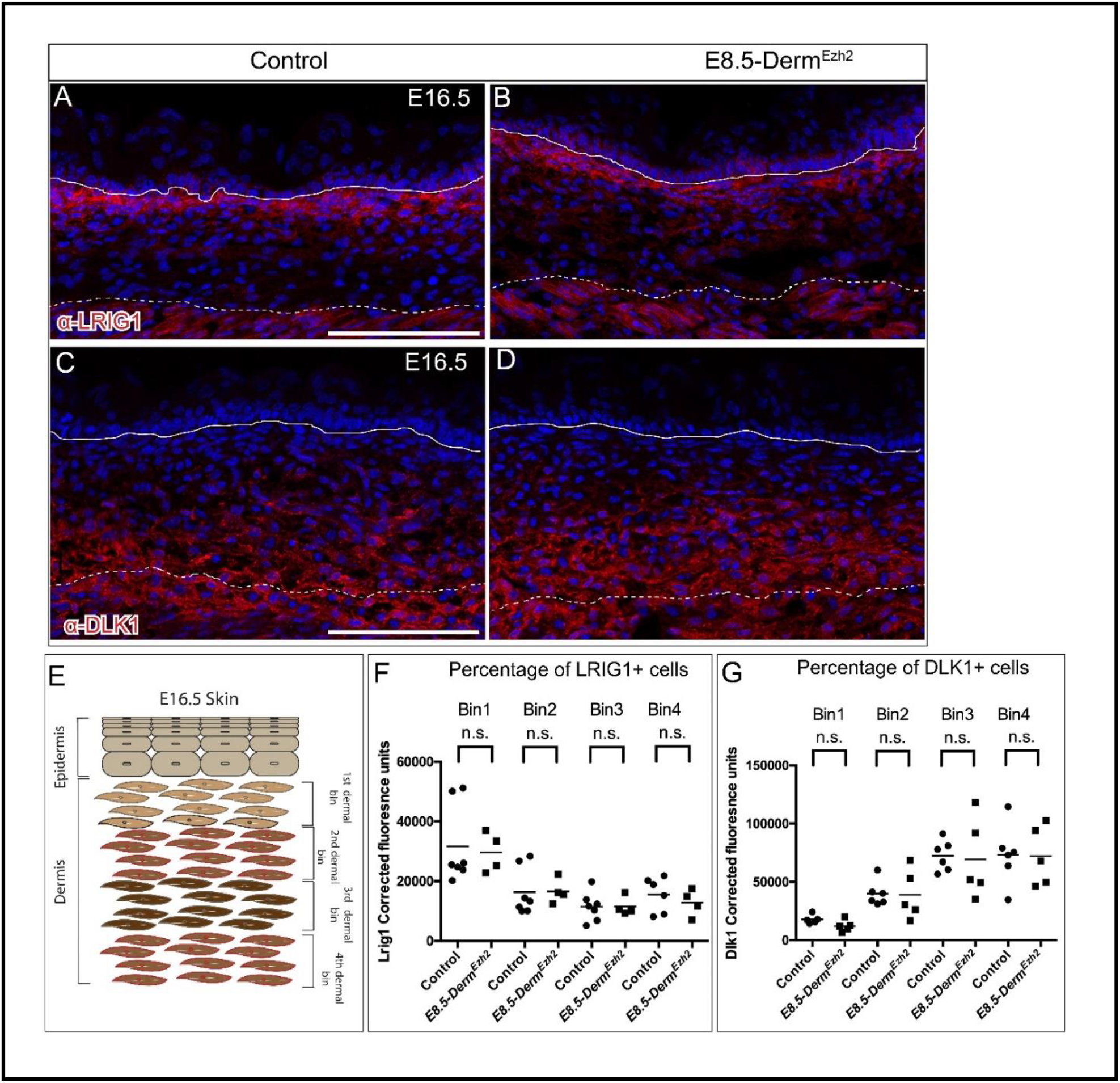
The PP marker, LRIG1, and the RP marker, DLK1, are comparable between controls and *E8.5-Derm*^*Ezh2*^ mutants: Indirect immunofluorescence with DAPI counterstain (A-D) showing papillary progenitor marker, LRIG1 (A, B), and reticular progenitor marker, DLK1 (C, D). Schematic illustration of binning of dermis employed to quantify LRIG1+/DLK1+ cells (E). Corrected fluorescence of LRIG1 (F) and DLK1 (G). Solid line demarcates the epidermis from dermis and dashed line demarcates the lower limit of the dermis. Scale Bar= 100 microns in A-D.

**FigureS4:**
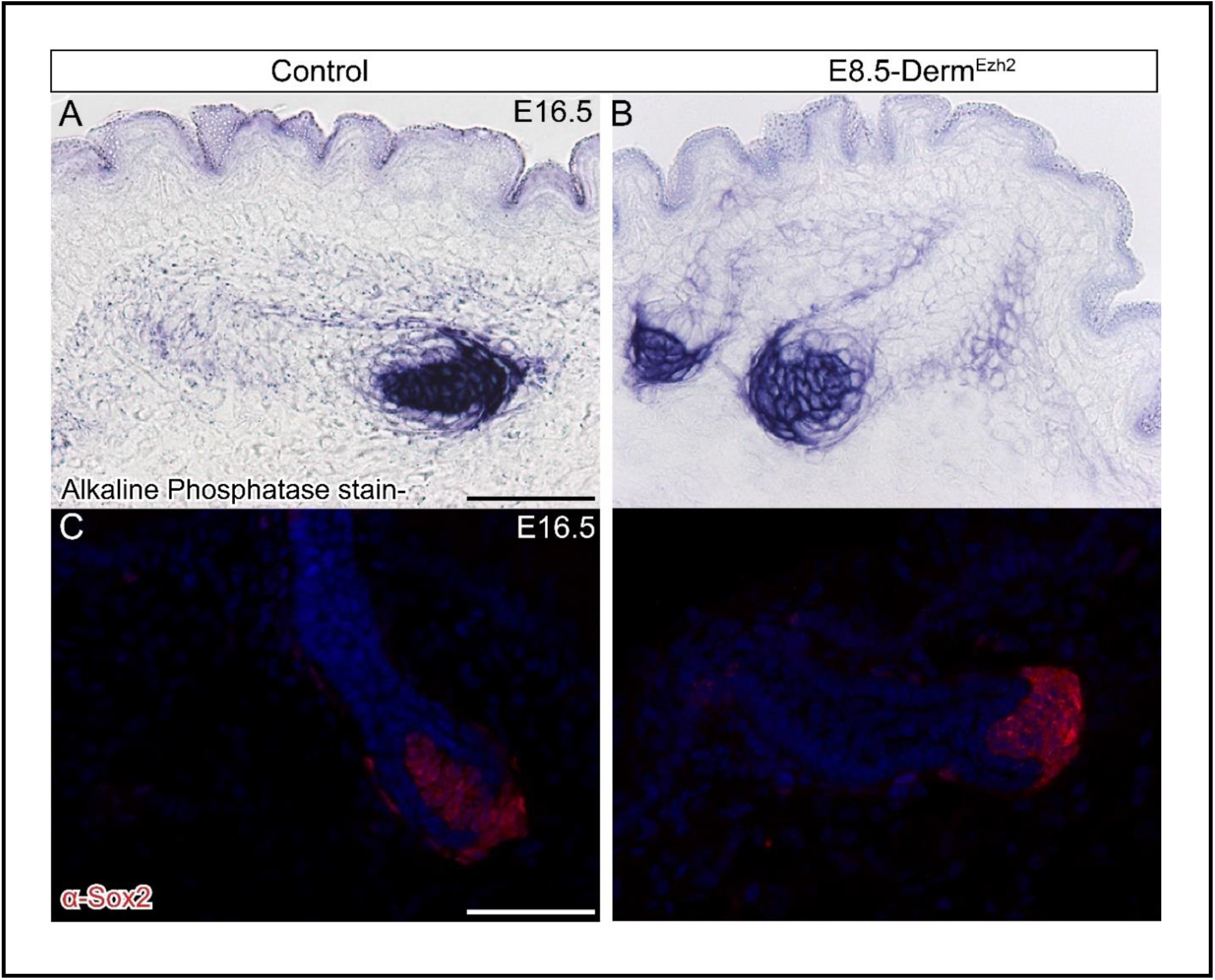
The dermal papilla marker expression is comparable between the controls and *E8.5-Derm*^*Ezh2*^ mice at E16.5. Alkaline phosphatase, a dermal papilla marker, staining sat E16.5 (A, B). Indirect immunofluorescence of dermal papilla marker, Sox2, with DAPI counterstain at E16.5 (C, D). Scale bar = 50 microns in A-D.

**FigureS5:**
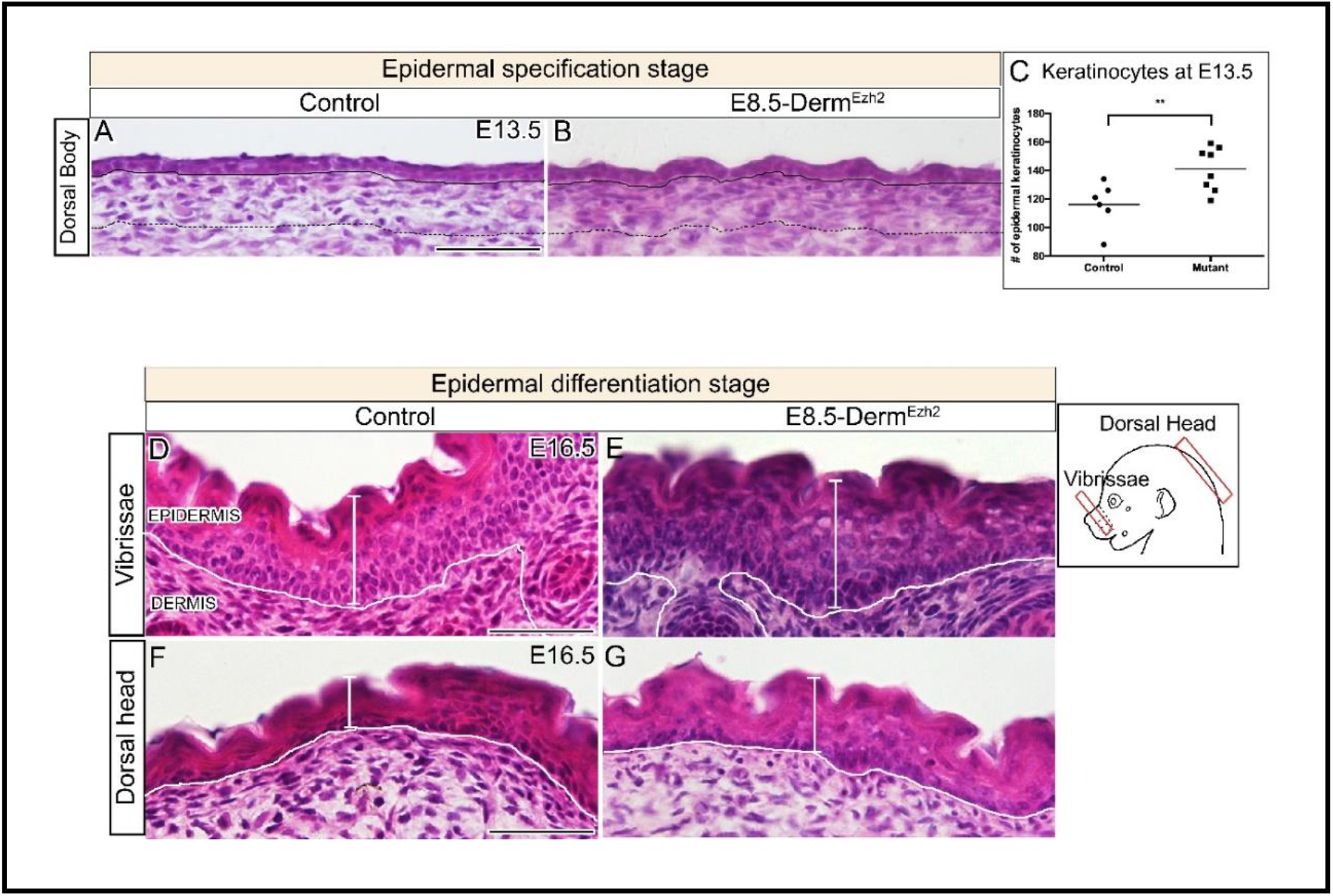
Epidermal hyperplasia in *E8.5-Derm*^*Ezh2*^ mice is evident at various locations and from E13.5 onwards: Hematoxylin and eosin staining of dorsal skin (A, B, D-G) showing the histology of the epidermis at E13.5 (A, B) and E16.5 (D-G). Quantification of the number of epidermal keratinocytes at E13.5 between controls and E8.5-Derm^Ezh2^ (C). Solid line demarcates the epidermis from dermis and dashed line demarcates the lower limit of the dermis. Scale Bar= 100 microns in A, B, D-G.

**FigureS6:**
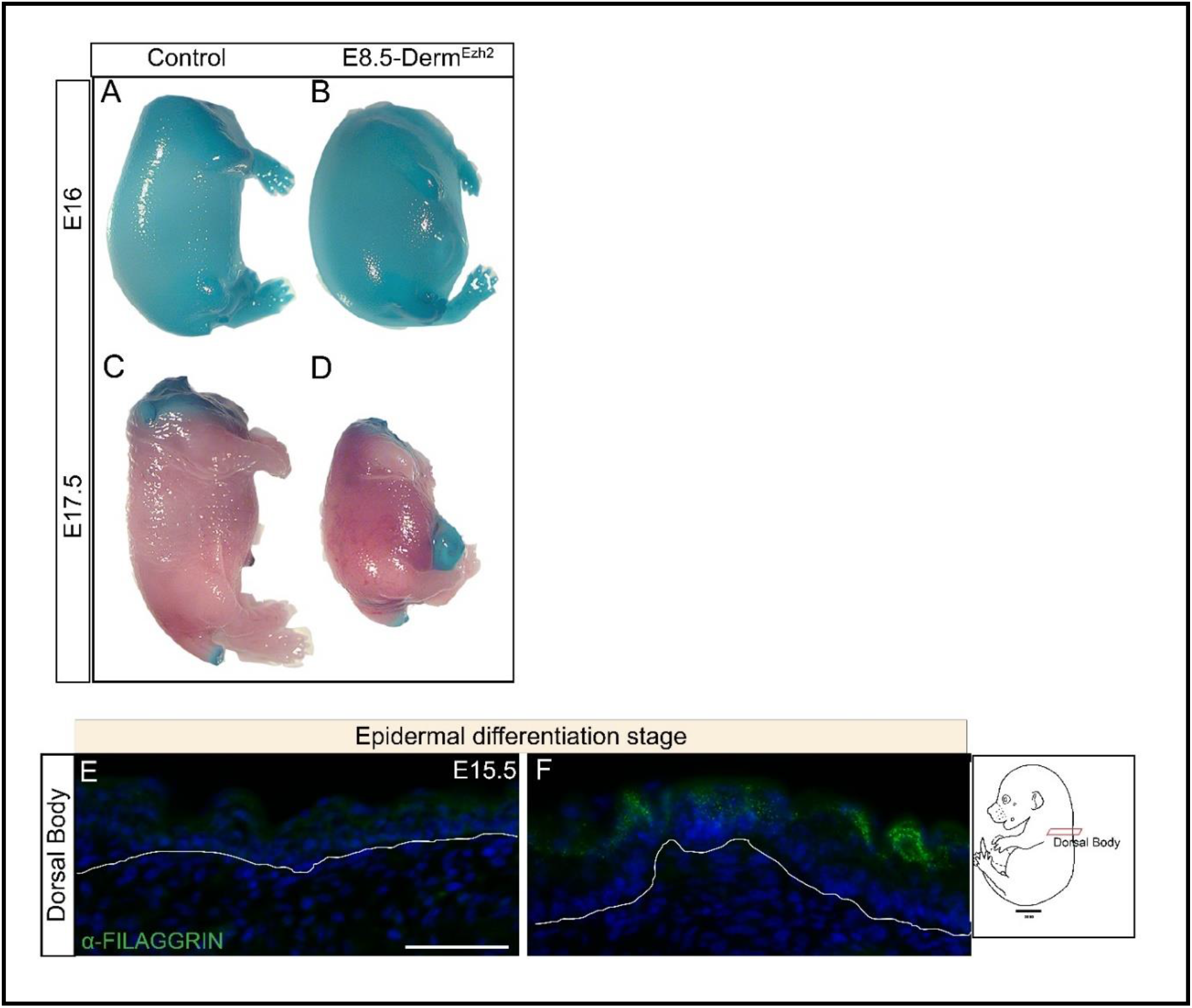
*E8.5-Derm*^*Ezh2*^ mice have intact barrier but exhibits accelerated differentiation: Whole mount toluidine blue O staining at E16 (A, B) and E17.5 (C, D). Indirect immunofluorescence of FILAGGRIN (E, F) with DAPI counterstain at E15.5. Solid line demarcates the epidermis from the dermis. Scale bar = 50 microns in A-D.

**Figure S7:**
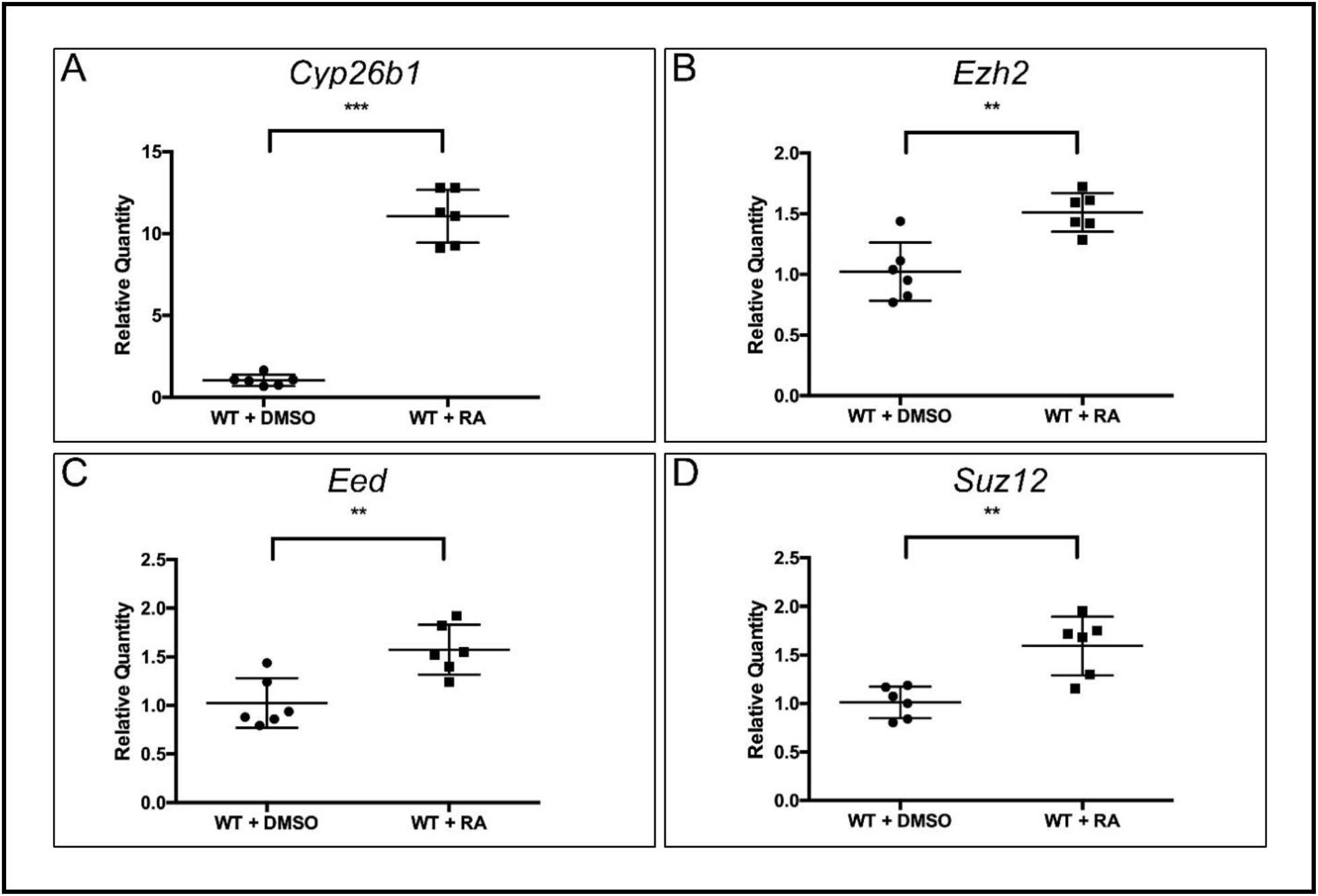
PRC2 components are not regulated by retinoic acid signaling. The relative quantity of *Cyp26b1* (A), a target of RA signaling, and PRC2 components (B-D) in RA treated wild-type skin in comparison to vehicle control treated embryos at E10.5.

